# Glycoprotein 130 Antagonism Counteracts Metabolic and Inflammatory Alterations to Enhance Right Ventricle Function in Pulmonary Artery Banded Pigs

**DOI:** 10.1101/2025.01.20.633954

**Authors:** Jenna B. Mendelson, Jacob D. Sternbach, Ryan A. Moon, Lynn M. Hartweck, Sophia R. Clark, Walt Tollison, Matthew T. Lahti, John P. Carney, Todd Markowski, LeeAnn Higgins, Felipe Kazmirczak, Kurt W. Prins

## Abstract

**Background:** Right ventricular dysfunction (RVD) is a risk factor for death in multiple cardiovascular diseases, but RV-enhancing therapies are lacking. Inhibition of glycoprotein-130 (GP130) signaling with the small molecule SC144 improves RV function in rodent RVD via anti-inflammatory and metabolic mechanisms. However, SC144’s efficacy and molecular effects in a translational large animal model of RVD are unknown.

**Methods:** 4-week-old castrated male pigs underwent pulmonary artery banding (PAB). After 3 weeks, PAB pigs were randomized into 2 groups (daily injections of SC144 [2.2 mg/kg, PAB-SC144, *n*=5] or vehicle [PAB-Veh, *n*=5] for 3 weeks). Five age-matched pigs served as controls. Cardiac MRI quantified RV size/function. Right heart catheterization evaluated hemodynamics. Single-nucleus RNA sequencing delineated cell-type specific changes between experimental groups. Electron microscopy evaluated RV mitochondrial morphology. Phosphoproteomics identified dysregulated RV kinases. Lipidomics and metabolomics quantified lipid species and metabolites in RV tissue. Quantitative proteomics examined RV mitochondrial protein regulation.

**Results:** SC144 significantly improved RV ejection fraction (Control: 60±4%, PAB-Veh: 22±10%, PAB-SC144: 37±6%) despite similar RV afterload. Single-nucleus RNA sequencing demonstrated PAB-Veh pigs had lower cardiomyocyte and higher macrophage/lymphocyte/pericyte/endothelial cell abundances as compared to control, and many of these changes were blunted by SC144. SC144 combatted the downregulation of cardiomyocyte metabolic genes induced by PAB. Kinome enrichment analysis suggested SC144 counteracted RV mTORC1 activation. Correspondingly, SC144 rebalanced RV autophagy pathway proteins and improved mitochondrial morphology. Integrated lipidomics, metabolomics, and proteomics analyses revealed SC144 restored fatty acid metabolism. Finally, CellChat analysis revealed SC144 restored pericyte-endothelial cell cross-talk.

**Conclusion:** GP130 antagonism blunts elevated immune cell abundance, reduces pro-inflammatory gene transcription in macrophages and lymphocytes, rebalances autophagy and preserves fatty acid metabolism in cardiomyocytes, and restores endothelial cell and pericyte communication to improve RV function.

## Introduction

Right ventricular dysfunction (RVD) is a risk factor for death in numerous cardiovascular diseases including systolic and diastolic heart failure^1^, pulmonary hypertension^2, 3^, congenital heart disease^4, 5^, and valvular cardiomyopathy^6^. In all of these conditions, RVD is predominately driven by heightened afterload, and at present, there are no therapeutic interventions that effectively combat the pathogenic processes in RVD. This is in direct opposition to the multiple therapies that improve survival in patients with left ventricular systolic dysfunction^7^. Our current inability to effectively treat RVD is further hampered by both the relative lack of understanding of the mechanisms of RVD, and evaluations of therapeutic approaches in translational large animal models of RVD^7^.

Glycoprotein-130 (GP130) is the master membrane receptor for the interleukin-6 (IL6) superfamily of inflammatory cytokines^8^, and GP130 signaling modulates many aspects of cellular biology including inflammation and metabolism in diverse cell types^8^. We nominated GP130 signaling as a target for RVD as GP130 antagonism with the small molecule SC144 improves RV function via restoration of mitochondrial morphology and metabolic function in the monocrotaline rat model of pulmonary hypertension-mediated RVD^9^. In addition, clinical data demonstrate GP130 signaling is associated with RVD in pressure overload conditions. Specifically, serum IL6 levels are independently associated with RVD in patients with pulmonary arterial hypertension (PAH)^10^. Furthermore, two independent studies show the IL6-JAK-STAT (janus kinase-signal transducer and activator of transcription) pathway is enriched in the RV transcriptome of patients with decompensated RVD due to PAH^11, 12^. Thus, the summation of preclinical and clinical data suggests antagonizing GP130 signaling may have therapeutic potential for RVD.

Mechanistically, GP130 could regulate RV function via mitochondrial metabolism through mammalian target of rapamycin complex 1 (mTORC1) signaling. GP130 activates mTORC1 signaling in primary mouse neonatal cardiomyocytes^13^. In addition, GP130 signaling via IL11 modulates systemic metabolism and mTORC1 signaling in aging^14^. In fact, antagonism of GP130 signaling via an IL11 blocking antibody counteracts metabolic derangements in several tissues in rodents, which ultimately extends lifespan. One way that mTORC1 regulates metabolism is suppression of autophagy^15^. Interestingly, GP130 signaling is already implicated in autophagy homeostasis^16, 17^. Autophagic flux is impaired in cardiac dysfunction, and this contributes to metabolic derangements in cardiomyocytes via the accumulation of dysfunctional mitochondria^15^. At present, the impacts of GP130 antagonism on autophagy, mitochondrial homeostasis, and metabolism in large animal RVD are unexplored.

Here, we evaluate the therapeutic utility of GP130 antagonism by determining how SC144 impacts right heart structure/function in pulmonary artery banded pigs. We integrate cardiac MRI and invasive hemodynamic evaluations of RV function with histological analyses, single-nucleus RNA sequencing, mitochondrial proteomics, electron microscopy, phosphoproteomics, lipidomics, and metabolomics to systemically define how GP130 signaling modulates right ventricular biology in a highly translational porcine model of RVD.

## Materials and Methods

### Pulmonary Artery Banding (PAB)

Ten Yorkshire castrated male pigs aged 20-40 days old and weighing 6.3±1 kg on the day of surgery were used in this study. Five castrated pigs of a similar age and weight were housed at the University of Minnesota Research Animal Resources Facility for six weeks, and these animals served as controls. Pulmonary artery banding was performed as previously described^18–20^.

### Animal Treatment Protocol

The small molecule SC144 was used to inhibit GP130 signaling, and vehicle control was propylene glycol (65%), DMSO (5%), and phosphate buffered saline (30%)^9, 21^. Daily SC144 or vehicle administration (2.2 mg/kg) began 3 weeks after pulmonary artery banding. We calculated the dose of SC144 by converting the effective dose from rodents to pigs using body surface area conversion^22^.

### Cardiac MRI

Cardiac MRI (cMRI) studies were performed with a Siemens Aera 1.5T scanner as we previously reported^18, 20^. These data were analyzed by an experienced cMRI cardiologist blinded to the treatment groups using standard software (Precession by Heart Imaging Technologies, Durham, NC). LV and RV end-diastolic and end-systolic volumes, ejection fractions and mass were quantified by planimetry of the end-diastolic and end-systolic endocardial and epicardial borders on the short-axis cine images. PAB stenosis percentage was performed by double orthogonal planes in the pulmonary artery to measure the area proximally and at the narrowest area of the PAB.

### Terminal data collection

After six weeks, animals were anesthetized to undergo right heart catheterization and a terminal cardiac MRI study. Venous blood was collected, and a Swan-Ganz catheter (Edwards Lifesciences) advanced through the right heart into the pulmonary artery to measure RV pressure and cardiac output. Upon completion of the scan, animals were humanely euthanized for tissue collection. This study was approved by the University of Minnesota Institutional Animal Care and Use Committee (IACUC).

### Cardiac histology

RV sections were fixed in 10% formalin, embedded in paraffin, and sectioned at 10 µm. Sections were stained with wheat germ agglutinin, and mounted with Prolong Glass Antifade Mountant containing NucBlue Stain (Hoechst 33342, Thermo Fisher Scientific). Images obtained on a Zeiss LSM900 Airyscan 2 confocal microscope, and cardiomyocyte cross-sectional area assessed with FIJI. To assess fibrosis, sections were stained with picrosirius red, images acquired on a Zeiss AxioCam IC, and analyzed using FIJI. Image collection and analysis was completed by scientists blinded to the treatment groups.

### Nuclei Isolation

Nuclei were isolated from four pigs per experimental group following an established protocol for frozen cardiac tissue^23^. Briefly, flash frozen tissue was homogenized in Nuclei EZ lysis buffer (Sigma Aldrich), filtered, and centrifuged to remove large debris. Nuclei suspensions were cleaned twice using density gradient centrifugations. Isolated nuclei were further purified using fluorescence activated cell sorting (FACS) after propidium iodide staining at the University of Minnesota Flow Cytometry Resource. All solutions were supplemented with RNAse inhibitor (0.2 U/µl, Millipore Sigma). Sequencing, library preparation and alignment to the pig genome (Sscrofa10.2) was completed by the University of Minnesota Genomics Center.

### Single nucleus RNAsequencing (SnRNASeq) analysis

SnRNASeq analysis was performed using Rstudio v4.4 accessed through the Minnesota Supercomputing Institute. Matrix files from CellRanger were transformed into Seurat objects and processed using Seurat v5^24^. Nuclei containing between 200-2500 genes (200-2500) and <5% mitochondrial DNA were used for downstream analysis. Potential doublets were identified and removed from each sample with doubletFinder^25^. Resulting Seurat objects were then combined for normalization, feature selection, scaling, dimensionality reduction, and clustering following the standard Seurat workflow^24^. Azimuth^26–29^ was used to guide clustering resolution to ensure different cell types clustered independently while preventing over resolution. We initially identified 27 clusters and then determined cell types at the cluster level. Cell type IDs were assigned by identifying the highly expressed genes in each cluster using the FindConservedMarkers function in Seurat. We cross-validated our cellular identifications with Single Cell Portal^30^, a database of previously established cell marker genes^31^, for up to 20 of the most highly expressed genes in each cluster to determine cell types. Three clusters did not contain genes highly expressed in any cell type and were therefore deemed to be noise and removed from subsequent analysis. Cell identity was confirmed using marker genes from several additional studies^26, 28, 29^. After cell types were identified, nuclei were pseudobulked for differential gene expression analysis using DESeq2^32^. Genes were considered differentially expressed if |log2FoldChange| ≥ 0.5 and adjusted *p*-value <0.05. Genes unique to cardiomyocytes that were present in non-cardiomyocyte cell types were removed by clearing atrial natriuretic peptide (NPPA), natriuretic peptide B (NPPB), and the top 100 expressed cardiomyocyte genes^28^.

We then performed a pathway analysis using ShinyGO, and pathways are shown if two or more pathways were enriched. For several comparisons, outlined here, *k*-means clustering was performed before pathway analysis using STRING v12.0^33^ When evaluating cardiomyocyte gene expression in PAB-Veh vs control and PAB-SC144 vs control, more than three unconnected groups were identified during clustering (*k*=3), and most genes were in cluster 1, which was then selected for clustering again (*k*=3). Endothelial cell DEGs comparing PAB-Veh and control were clustered first, and cluster 1 was used for the pathway analysis. Pericyte, lymphocyte, and macrophage pathway analysis was performed without clustering. Correlation analysis of nuclei relative abundance and ejection fraction was completed with Metaboanalyst. The number of cardiomyocyte subclusters was identified during initial cell clustering. Genes that distinguished cardiomyocyte subclusters were identified using the “FindConservedMarkers” function in Seurat and comparing expression in each subcluster to all other cardiomyocyte subclusters. The top 250 genes were clustered, and cluster 1 was used for pathway analysis. The CellChat v2 package evaluated predicted cell-cell communication, following their standard workflow^34^.

### Phosphoproteomics Analysis

Flash frozen RV samples were enriched for phosphorylated peptides and analyzed by mass spectrometry at the University of Minnesota Center for Metabolomics and Proteomics^35^. Sequenced proteins that did not have identified phosphorylation sites were removed. Proteins that were significantly (adjusted *p*-value<0.05, Proteome Discoverer Software) elevated or reduced in PAB-Veh or PAB-SC144 compared to control were selected for kinome enrichment analysis using Kinase Enrichment Analysis v3^36^. The top 10 kinases identified by MeanRank score using Kinase Enrichment Analysis were visualized on kinome maps using Coral^37^

### RV Mitochondrial Proteomics

RV mitochondrial enrichments were isolated from frozen RV tissue from all 15 pigs used in this study with a mitochondrial isolation kit (Abcam, Cambridge, MA) as previously described for TMT 16-plex proteomics analysis at the University of Minnesota Center for Metabolomics and Proteomics^9, 18, 38^.

### iPSC-CM differentiation

Human induced pluripotent stem cells (iPSC, WTC, Allen Institute) tagged with a green fluorescent protein to the translocase of outer mitochondrial membrane 20 (TOMM20) were differentiated into cardiomyocytes following Allen Institute protocols^39^. Briefly, cells were seeded (100,000 cells/well) onto Matrigel (337 µg/mL, Corning) coated 12-well dish. At 70-85% confluency cells were treated with CHIR99021 (7.5 µmol) in Roswell Park Memorial Institute (RPMI) media supplemented with B27- (RPMI/B27-). 48 hours later cell media was replaced with RPMI/B27- plus IWP (7.5 µmol). After 48 hours media was changed to RPMI/B27-. 48 hours later media was changed to RMPI supplemented with B27+ (RPMI/B27+). Media was subsequently changed every 48 hours with fresh RMPI/B27+ until cells were beating, then media change frequency was reduced to twice per week.

### iPSC-CM in-vitro experiments

Differentiated iPSC-CMs were passaged onto Matrigel coated coverslips in RMPI/B27+ media with rho kinase (ROCK) inhibitor (10 µm). 24 hours after passaging cell media was replaced with RPMI/B27+. The following day, cells were treated overnight with either the GP130 ligand oncostatin M (OSM, 200 ng/mL, R&D Systems), or the autophagy inhibitor Bafilomycin (BAF, 50 nmol, Selleck Chemicals). Vehicle control was phosphate buffered saline. In a separate experiment, iPSC-CMs were given SC144 (10 µmol, Selleck Chemicals) plus either OSM or BAF overnight.

### Electron microscopy

RV free wall samples (1-2mm^3^) were fixed in 4% paraformaldehyde, 1% glutaraldehyde in 0.1M phosphate buffer, pH 7.2 for electron microscopy. Tissue was sliced into ultrathin (0.1 µM) sections, and micrographs acquired using a JEOL 1400 Plus transmission electron microscope at 80 kV with a Gatan Orius camera at the Mayo Clinic. Mitochondrial length to width ratio was quantified using FIJI to assess morphology. Cristae structure was evaluated by applying a cristae score based on the percentage of each mitochondrion occupied by cristae^40^. Measurements were completed by scientists blinded to the treatment group.

### RV Lipidomics and Metabolomics

Lipidomic and metabolomic profiling of flash frozen RV tissue was performed by Metabolon Inc. (Durham, NC) as formerly reported^9, 41^. The Metabolon platform used ultrahigh performance liquid chromatography-tandem mass spectroscopy on a Waters ACQUITY ultra-performance liquid chromatography and a Thermo Scientific Q-Exactive high resolution/accurate mass spectrometer interfaced with a heated electrospray ionization source and Orbitrap mass analyzer. Batch-normalized data were used for downstream analysis.

### Integrative Bioinformatics Analysis and Statistics

Global changes in proteomics, lipidomics, and metabolomics data were assessed using hierarchical cluster analysis, partial least squares discriminant analysis (PLS-DA), and random forest classification using Metaboanalyst^42^. The top 250 proteins identified by PLS-DA were selected for a pathway analysis using the Kyoto Encyclopedia of Genes and Genomes (KEGG) database in ShinyGO v0.81^43^. Lipidomic pathway analysis was completed using Lipid Ontology (LION)^44^. The top 200 metabolites identified by hierarchical cluster analysis were selected for a pathway analysis using Metaboanalyst.

### Western blots

Protein was isolated from flash-frozen RV tissue from each pig was extracted in sodium dodecyl sulfate (SDS) buffer. 25 μg of protein extracts were used for immunoblotting using the Odyssey Infrared Imaging system as previously described^9, 38^. Post-transfer SDS-PAGE gels were stained with Coomassie brilliant blue (CBB) and the band corresponding to myosin heavy chain quantified and used as the loading control.

### Statistical Analysis

Data normality was evaluated using Shapiro-Wilk test. To compare the means of three groups, one-way analysis of variance (ANOVA) with Tukey post-hoc analysis was used when the data were normally distributed. If the data were not normally distributed, Kruskal-Wallis ANOVA with Dunn’s post-hoc analysis was used. When comparing the means of two groups, unpaired t-test was used if data were normally distributed. If the data were not normally distributed, the Mann-Whitney U-test was completed. Statistical significance was defined as *p*-value <0.05. Statistical analyses and graphing were performed with GraphPad Prism version 10. Data are presented as mean ± standard deviation if normally distributed or median and interquartile range if not normally distributed. Graphs show the mean or median and all individual values.

### Figure Preparation

Figures were prepared in Adobe Illustrator 2024.

## Results

### GP130 antagonism enhanced RV function and counteracted RV cardiomyocyte hypertrophy

First, we evaluated the effects of SC144 treatment on cardiac anatomy and function in control (*n*=5), pulmonary artery banded (PAB) pigs treated with vehicle (PAB-Veh, *n*=5), and PAB pigs treated with SC144 (PAB-SC144, *n*=5). Blinded cardiac magnetic resonance imaging (cMRI, **Figure 1A, Supplemental Table 1**) analysis revealed SC144 significantly enhanced ejection fraction (RVEF) as compared to PAB-Veh pigs (control: 60±4%, PAB-Veh: 22±10%, PAB-SC144: 37±6%, Figure 1B). PAB-Veh and PAB-SC144 pigs had comparable RV afterloads determined by RV systolic pressure divided by cardiac output (Figure 1C) as RV systolic pressure relative to cardiac output and magnetic resonance angiography determined PA banding severity was equivalent between the two groups (Figure 1D), which suggested SC144 had a positive inotropic effect on the RV. Consistent with this, there were nonsignificant improvements in right atrial size and function in PAB-SC144 pigs. (**Supplemental Table 1**). Next, we evaluated how SC144 impacted RV histology and found it prevented cardiomyocyte hypertrophy (Figure 1E-F, control: 131±54 µm^2^, PAB-Veh 200±70 µm^2^, PAB-SC144: 129±62 µm^2^), but did not significantly change fibrosis (**Supplemental Figure 1**).

**Figure 1:**
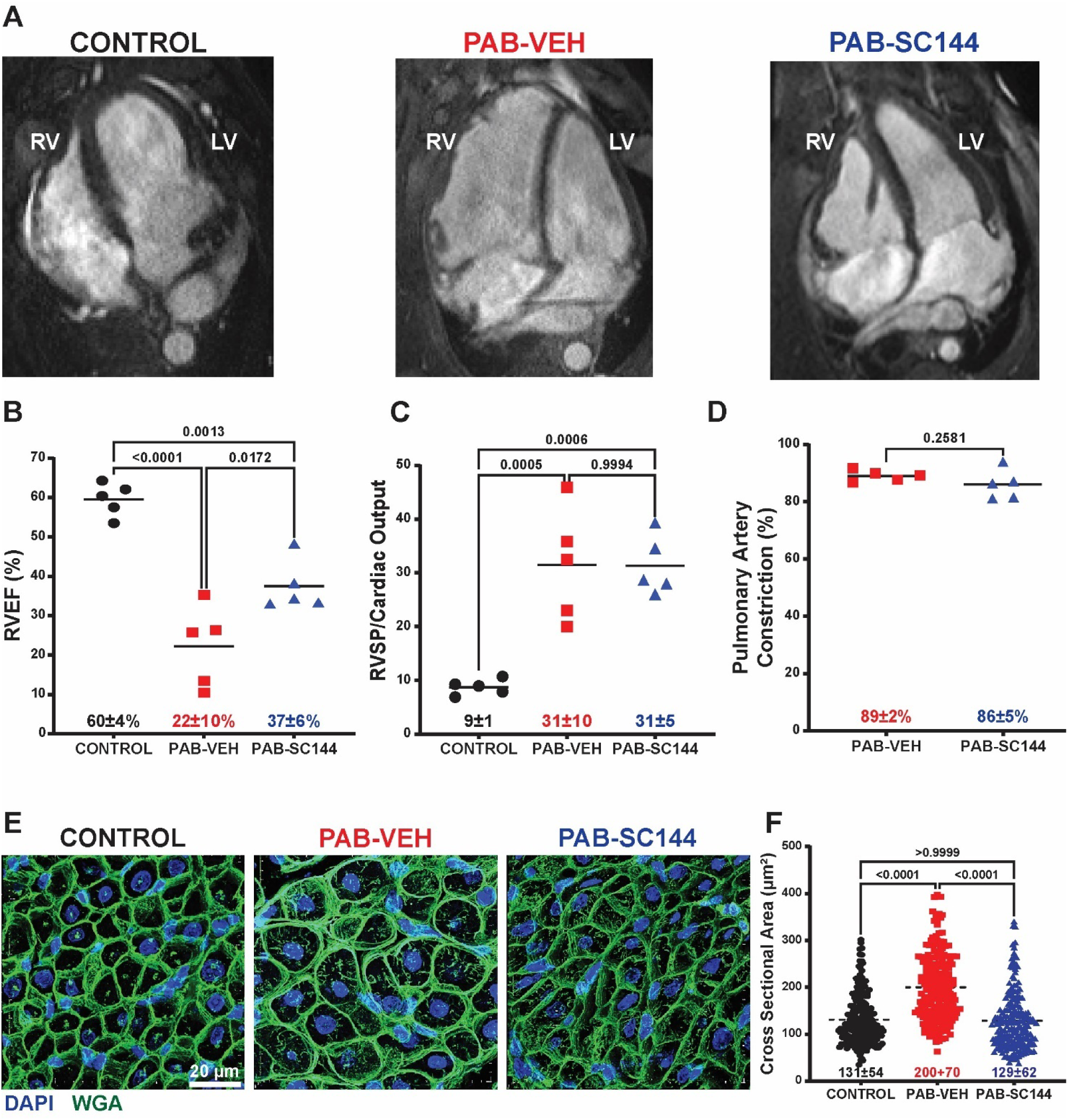
GP130 antagonism enhanced RV function and mitigated cardiomyocyte hypertrophy. (**A**) Representative still images from terminal cMRI studies. (**B**) Treatment with SC144 significantly enhanced RV function compared to PAB-Veh pigs (control: 60±4%, PAB-Veh: 22±10%, PAB-SC144: 37±6%) (**C**) Afterload was similar between PAB-Veh and PAB-SC144 pigs (PAB-Veh: 31±10, PAB-SC144: 31±5). (**D**) Pulmonary artery constriction was comparable in PAB-Veh and PAB-SC144 pigs (PAB-Veh: 89±2%, PAB-SC144: 86±5%). **(E**) Representative confocal microscopy images of RV sections stained with wheat germ agglutinin (green: WGA, blue: DAPI) to evaluate cardiomyocyte hypertrophy. (**F**) Treatment with SC144 prevented cardiomyocyte hypertrophy (control: 131±54 µm^2^, PAB-Veh: 200±70 µm^2^, PAB-SC144: 129±62 µm^2^). Kruskal-Wallis ANOVA with Dunn post hoc test in **F**.

### snRNA sequencing defined changes in the RV cellular landscape and nominated cell types associated with RV function

To investigate cell-specific alterations caused by PAB and treatment with SC144, we performed single-nucleus RNA sequencing (snRNAseq) from RV free wall sections of four pigs from each experimental group (Figure 2A). Unsupervised clustering using Seurat^24^ identified 8 cell types: macrophages, fibroblasts, lymphocytes, endothelial cells, pericytes, smooth muscle cells, cardiomyocytes, and neuronal cells (**Figure 2B-C**). The cell composition of the RV was distinct in the three experimental groups, and the relative abundance of cardiomyocytes (control: 70.3%, PAB-Veh: 51.2%, PAB-SC144: 59.3%), macrophages (control: 6.2%, PAB-Veh: 13.3%, PAB-SC144: 11.0%), lymphocytes (control: 2.8%, PAB-Veh: 8.2%, PAB-SC144: 5.0%), pericytes (control: 3.9%, PAB-Veh: 7.3%, PAB-SC144: 6.5%), and endothelial cells (control: 2.4%, PAB-Veh: 4.2%, PAB-SC144: 5.0%) were the most discrepant between the three groups (Figure 2D-E). The reduction in cardiomyocytes and the increase in macrophage and lymphocyte relative abundances in PAB-Veh were partially blunted by treatment with SC144 (Figure 2F).

**Figure 2:**
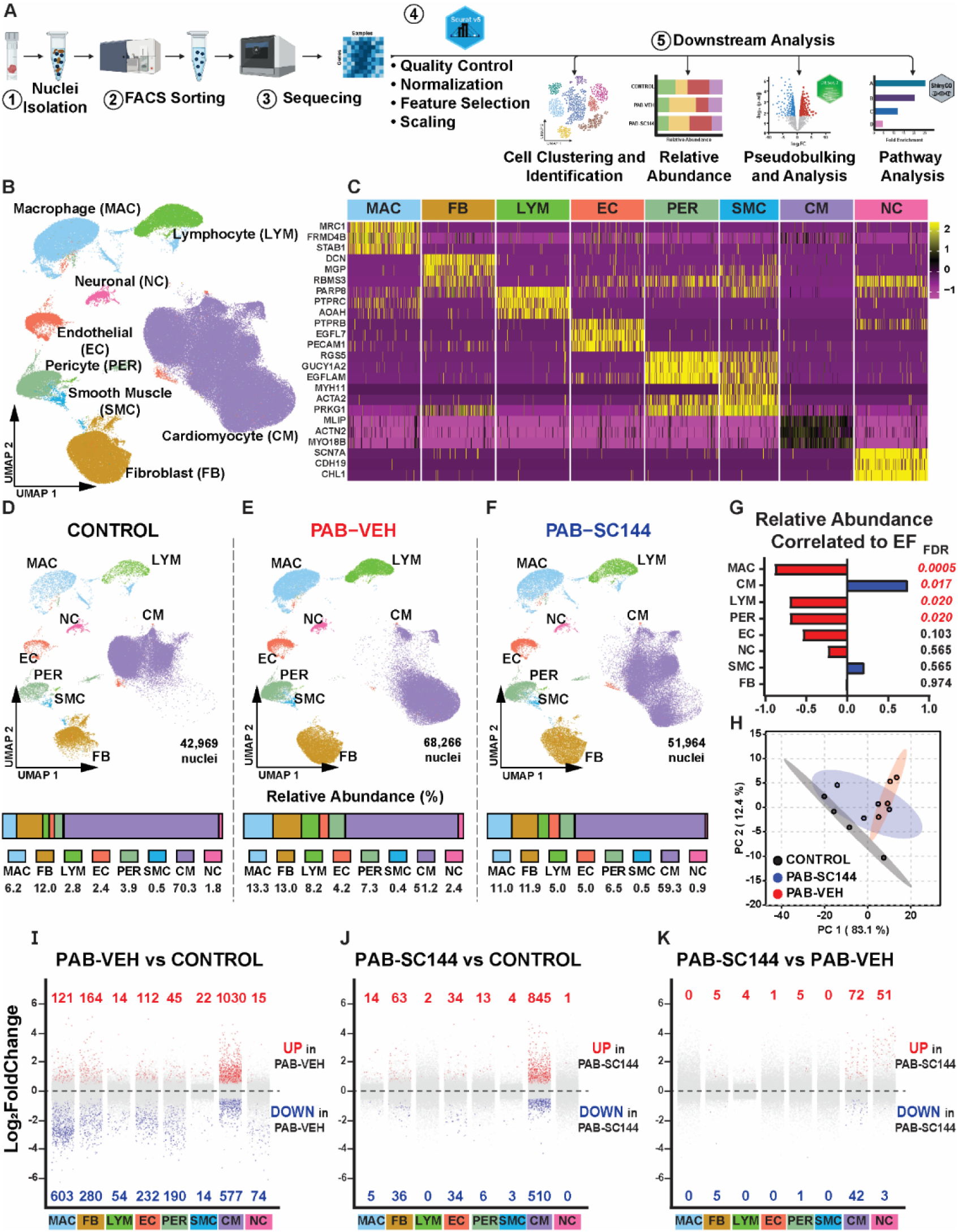
snRNAseq identified alterations in the cellular landscape in the three experimental groups. (**A**) Steps of single-nucleus RNA sequencing (snRNAseq) analysis. (**B**) Uniform manifold approximation and projection (UMAP) visualization of 8 cell types identified with unsupervised clustering. (**C**) Canonical markers used to validate cell type identities. UMAP and relative abundances of each cell type in (**D**) control, (**E**) PAB-Veh, and (**F**) PAB-SC144 RVs. (**G**) Correlation analysis of relative abundance and ejection fraction (EF). The relative abundances of macrophages, lymphocytes, and pericytes were significantly negatively associated with EF, and cardiomyocyte abundance was significantly positively correlated to EF. (**H**) Principal component analysis separated experimental groups using nuclei relative abundance. Differential expression analysis of pseudobulked data from all 8 cell types (red: Log2FoldChange>0.5 and adjusted p-value<0.05, blue: Log2FoldChange<-0.5 and adjusted p-value<0.05) comparing (**I**) PAB-Veh vs control, (**J**) PAB-SC144 vs control, and (**K**) PAB-SC144 vs PAB-Veh. The comparison between PAB-Veh and control had the most DEGs, and cardiomyocytes had the most DEGs across comparisons.

Next, we performed a correlation analysis to define the relationship between RVEF and the relative abundances of each cell to determine which cell types were associated with RV function. Cardiomyocyte relative abundance was positively correlated with RVEF (Figure 2G), while the relative abundances of macrophages, lymphocytes, and pericytes were significantly and negatively associated with RVEF. To further probe cell type differences between experimental groups, we completed a principal component analysis (PCA) and found SC144 animals were an intermediate between control and PAB-Veh (Figure 2H).

### snRNAseq delineated alterations in gene regulation in distinct cell subtypes in the RV

Pseudobulking and differential gene expression analysis calculated the number of differentially expressed genes (DEGs) for each cell type comparing PAB-Veh and control (Figure 2I), PAB-SC144 and control (Figure 2J), and PAB-SC144 and PAB-Veh (Figure 2K). Of all the comparisons, the most divergent genetic programs were identified when analyzing PAB-Veh versus control. Macrophages and cardiomyocytes had the greatest number of DEGs when controls and PAB-Veh were compared. Gene expression between PAB-SC144 and PAB-Veh RVs was more similar. Again, cardiomyocytes had the largest number of DEGs and many cell types, including macrophages, had none. These data indicated that PAB most robustly altered the genetic landscape in cardiomyocytes and macrophages, and SC144 predominately modulated cardiomyocyte genetic regulation.

### SC144 reduced the abundance of stressed cardiomyocytes and counteracted metabolic gene expression dysregulation in RV cardiomyocytes

Because cardiomyocyte (CM) relative abundance was positively correlated with RVEF and CMs had the highest number of DEGs, we investigated CM in greater detail. Unsupervised clustering identified eight CM subgroups (Figure 3A), and the composition was distinct amongst the three experiment groups (Figure 3B-C).

**Figure 3:**
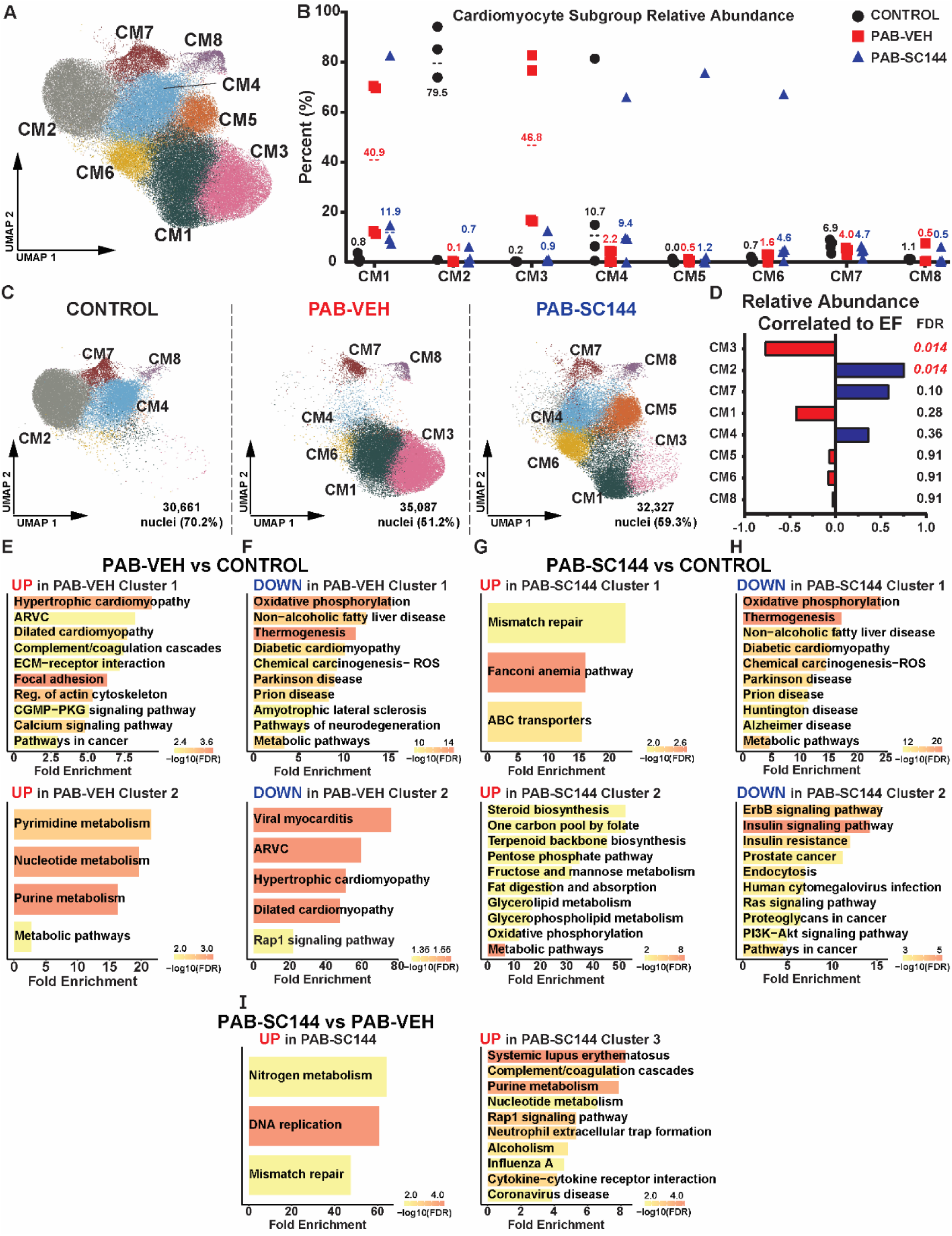
Cardiomyocyte metabolic genetic programming was altered by PAB but partially rescued with SC144 treatment. (**A**) UMAP displaying 8 subgroups of cardiomyocytes (CM). (**B**) Percent of CM nuclei from each pig across subgroups. (**C**) UMAP visualizing changes in cardiomyocyte subclusters between experimental groups. (**D**) Correlation analysis of relative abundance of cardiomyocyte subclusters and ejection fraction (EF). CM3 was negatively correlated to EF and CM2 was positively correlated to EF. Pathway analysis of DEGs (**E**) increased and (**F**) decreased in PAB-Veh nuclei compared to control. Pathways of DEGs (**G**) elevated and (**H**) decreased in PAB-SC144 compared to control. Metabolic pathways were altered by PAB and treatment with SC144, and downstream GP130 signaling was reduced in PAB-SC144 compared to control. (**I**) Pathway analysis of DEGs upregulated in PAB-SC144 compared to PAB-Veh. There were no enriched pathways comparing DEGs downregulated in PAB-SC144 and PAB-Veh.

The proportion of nuclei from subgroup 3 (CM3) was negatively correlated with RVEF, and subgroup 2 (CM2) was positively correlated with RVEF (Figure 3D). We then performed a pathway analysis of genes that subclustered each population to determine how PAB and SC144 modulated cardiomyocytes. We focused on CM3 because it was negatively correlated with RVEF and PAB-Veh RVs had the highest proportion of CM3, and treatment with SC144 reduced the abundance of this subtype (control: 0.2%, PAB-Veh: 46.8%, PAB-SC144: 0.9%). Five of the ten pathways in CM3 were associated with inflammation, and the mTOR signaling pathway was enriched in these cardiomyocytes (**Supplemental Figure 2A**). Then, we probed CM4 because treatment with SC144 increased the proportion of this subgroup to a similar level as control cardiomyocytes (control: 10.7%, PAB-Veh: 2.2%, PAB-SC144: 9.4%). Metabolic pathways, including oxidative phosphorylation were enriched in CM4 (**Supplemental Figure 2B**). Thus, these data suggested SC144 counteracted alterations in inflammation, metabolism, and mTOR in distinct RV cardiomyocyte subpopulations.

Finally, we performed pathway analysis of pseudobulked transcripts from DEGs between treatment groups to further evaluate how the total population of cardiomyocytes was altered. Multiple cardiomyopathy and nucleic acid metabolism pathways were elevated in PAB-Veh cardiomyocytes compared to control (Figure 3E). Pathways predicted to be downregulated in PAB-Veh compared to control were primarily cardiomyopathy and metabolic pathways (Figure 3F). Specifically, oxidative phosphorylation and lipid/fatty acid metabolism was suppressed in PAB-Veh cardiomyocytes. On the contrary, metabolic pathways, especially related to fat/lipid metabolism, were increased in PAB-SC144 cardiomyocytes compared to control (Figure 3G). Many of the pathways suppressed in PAB-SC144 compared to control were similar to those reduced in PAB-Veh (Figure 3H). However, phosphoinositide 3 kinase-protein kinase B (PI3K-AKT) and Ras signaling pathways, which are downstream of GP130, were reduced in PAB-SC144 treated cardiomyocytes compared to control. When comparing PAB-SC144 to PAB-Veh cardiomyocytes, DNA replication and repair pathways were enriched (Figure 3I). Overall, these data suggested RVD was marked by impaired cardiomyocyte metabolism via downregulation of oxidative phosphorylation and fatty acid metabolism. Treatment with SC144 suppressed GP130 gene activation in cardiomyocytes, and increased expression of fat/lipid metabolizing genes.

### SC144 exerted an anti-inflammatory effect

To further define the anti-inflammatory effect of SC144, we analyzed the genetic programs of macrophage and lymphocyte populations in all three experimental groups. In PAB-Veh macrophages, pathways that regulate macrophage differentiation, survival, and proliferation were suppressed when compared to control (**Supplemental Figure 3A**). The toll-like receptor signaling pathway, a component of the innate immune system that initiates pro-inflammatory gene transcription^45^, was enriched in PAB-Veh macrophages, which suggested a pro-inflammatory phenotype was present. The cyclic guanosine monophosphate-protein kinase G (cGMP-PKG) pathway was reduced in PAB-Veh lymphocytes compared to control (**Supplemental Figure 3B**). Because cGMP suppresses T cell activation^46^, the PAB-Veh lymphocytes were likely activated. Oxidative phosphorylation was increased in PAB-Veh lymphocytes relative to control, and because pro-inflammatory lymphocytes require heightened oxidative phosphorylation^47, 48^, which further supports the hypothesis that lymphocytes are pathogenically activated in the PAB-Veh RV. Interestingly, there were no DEGs comparing PAB-Veh to SC144 macrophages. In addition, we used CellChat^34^ to predict changes in cell-to-cell communication between experimental groups. PAB-Veh macrophages communicated more with cardiomyocytes, endothelial cells, and pericytes than control macrophages (**Supplemental Figure 3C**). Notably, SC144 suppressed cellular communication of macrophages to other cell types in the RV when compared to control or PAB-Veh (**Supplemental Figure 3D-3E**). The summation of these data implied SC144 combatted a pro-inflammatory RV milieu induced by both macrophages and lymphocytes.

### Phosphoproteomics demonstrated SC144 normalized predicted mTORC1 related signaling

To evaluate how SC144 impacted GP130 and potential off-target intracellular signaling in an unbiased manner, we performed a phosphoproteomics evaluation of RV free wall specimens and then kinase enrichment analysis of differentially expressed phosphoproteins (Figure 4A). In the PAB-Veh RV, two of the top ten identified kinases were mitogen activated protein kinases, supporting the hypothesis that GP130 signaling is activated in porcine RVD (Figure 4B). PAB was also predicted to have heightened ribosomal protein S6 kinase (RPS6KB1) activity, a mTORC1 subunit^49^. In contrast, many polo-like kinases (PLK), which activate mitophagy and antagonize mTORC1^50^, were enriched in PAB-SC144 RVs compared to control (Figure 4C). Furthermore, the abundance of phospho-mTOR was elevated in PAB-Veh compared to control and PAB-SC144 (**Supplemental Table 2**). Moreover, GP130 antagonism with SC144 depressed multiple pyruvate dehydrogenase kinases (PDKs, **Supplemental Figure 4A-B**), which support the utilization of glucose over fatty acid oxidation^51^. In particular, predicted PDK1 activity, which activates RPS6KB1^52^, was reduced in PAB-SC144 RVs compared to control. These data provided additional evidence that SC144 suppressed ectopic GP130 and mTORC1 signaling pathways.

**Figure 4:**
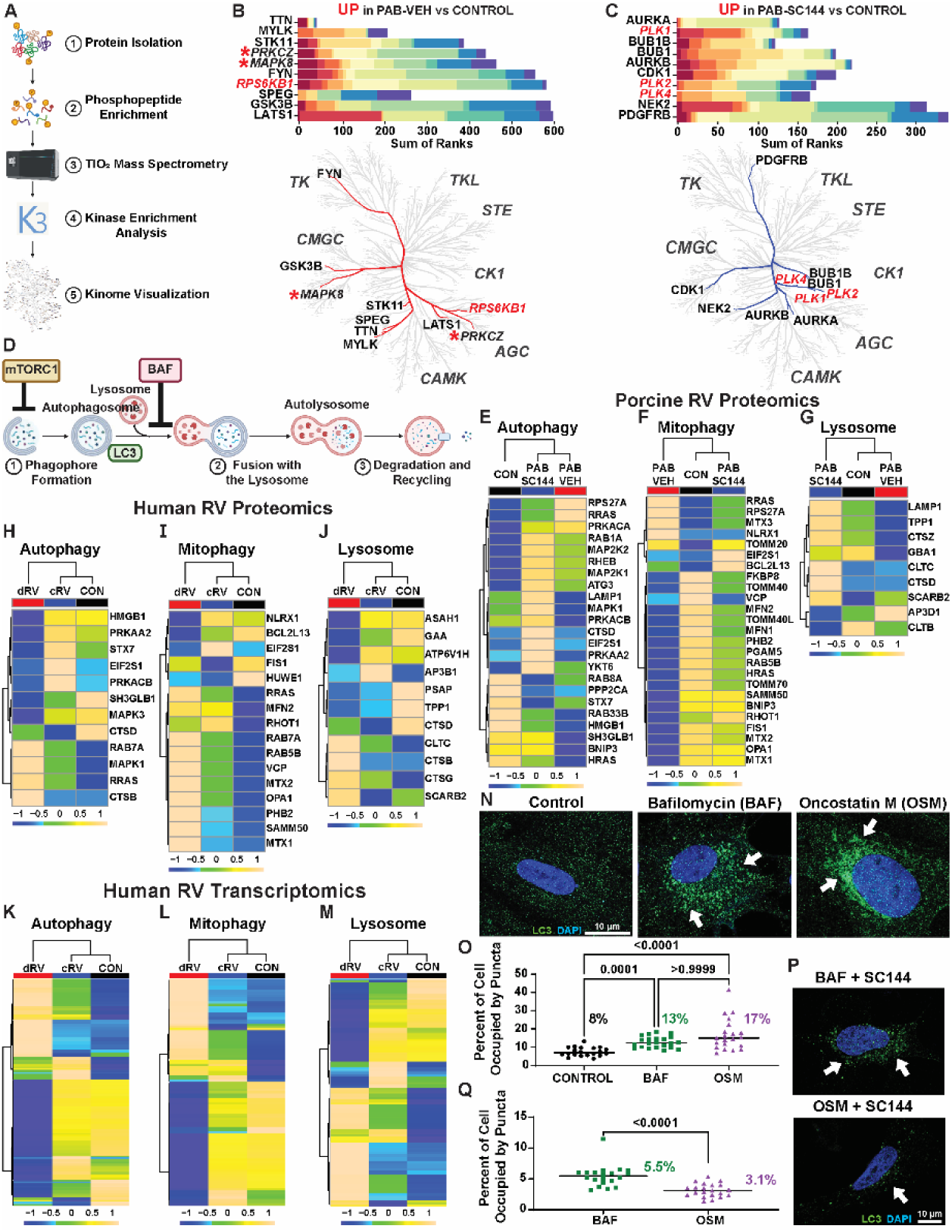
SC144 counteracted mTORC1-associated autophagic alterations. (**A**) Schematic of phosphoproteomics analysis. (**B**) Two (MAPK, PRKCZ, italicized, red stars) of the top 10 kinases identified in PAB-Veh were downstream of GP130. The mTORC1 subunit, RPS6KB1 (red italics) was also predicted to be enriched. (**C**) In PAB-SC144 RVs, several polo-like kinases (PLK, red italics), which antagonize mTORC1 were predicted to be enriched. (**D**) Primary steps in autophagy. Contents are isolated in an autophagosome, which fuses with a lysosome where the products are degraded and recycled. mTORC1 inhibits the initiation of autophagy. The autophagy inhibitor, bafilomycin (BAF), prevents lysosome and autophagosome fusion. Light chain 3 (LC3) is a component of autophagosomes. Porcine RV proteomics of the (**E**) autophagy, (**F**) mitophagy, and (**G**) lysosome pathways revealed many proteins were downregulated in PAB-Veh, but normalized with SC144 treatment. (**H**) Human PAH patients with decompensated RV function (dRV) had mostly decreased abundances of autophagy pathway proteins. (**I**) Mitophagy pathway proteins were mostly upregulated in dRV patients. (**J**) dRV patients had primarily decreased abundances of lysosome pathway proteins. Transcriptionally, similar numbers of genes had reduced expression between dRV, and compensated RV (cRV) patients in the (**K**) autophagy and (**L**) mitophagy pathway, but hierarchical cluster analysis clustered cRV and control groups together. (**M**) Lysosome pathway transcripts were mostly increased in cRV patients compared to dRV and control. (**N**) Representative confocal micrographs of human induced pluripotent stem cells differentiated into cardiomyocytes (iPSC-CMs) showed treatment with the autophagy inhibitor BAF or the GP130 ligand oncostatin M (OSM) increased LC3 positive puncta (white arrows: representative LC3 puncta; green: LC3; blue: DAPI) quantified as (**O**) the percent area occupied by LC3 puncta. (**P**) Representative confocal micrographs of subsequent iPSC-CMs treated with either BAF or OSM with SC144 demonstrated SC144 treatment prevented LC3 accumulation (white arrows, green: LC3, blue: DAPI) quantified as (**Q**) the percent area occupied by LC3. In **E**-**M** Scale bars show relative changes between groups.

### SC144 mitigated the reduction in autophagy/mitophagy/lysosomal proteins

Because mTORC1 signaling was enriched in PAB-Veh pigs, but not PAB-SC144 pigs in both phosphoproteomics and snRNAseq datasets, we performed mitochondrial proteomics and probed proteins in the autophagy, mitophagy, and lysosomal pathways (Figure 4D) to test the hypothesis that SC144 combatted pathogenic mTORC1-associated autophagy dysregulation. Hierarchical cluster analysis demonstrated most autophagy proteins were reduced in PAB-Veh pigs, but many were elevated in PAB-SC144 pigs as compared to controls (Figure 4E). Similarly, the majority of mitophagy proteins were downregulated in PAB-Veh when compared to controls, but treatment with SC144 normalized protein abundances (Figure 4F). Finally, nearly all proteins in the lysosome pathway were lowered in PAB-Veh RVs, but increased in PAB-SC144 RVs (Figure 4G).

### Autophagy pathway proteins and transcripts were downregulated in the dysfunctional RV of PAH patients

To examine the potential human relevance of our findings, we evaluated autophagy-related proteins and transcript changes in human PAH patients with RVD using publicly available proteomics and bulk RNA-sequencing data^11^. PAH patients were subdivided into compensated (cRV) and decompensated (dRV) RV function, and patients with dRV had mostly decreased abundances of autophagy pathway proteins compared to cRV and controls (Figure 4H). However, the proteins in the mitophagy pathway were mostly increased in dRV patients compared to cRV and healthy controls (Figure 4I). In the lysosome pathway, most proteins were downregulated in dRV patients, similar to the autophagy pathway (Figure 4J). Transcripts in the autophagy and mitophagy pathways had similar numbers of up and down regulated genes between patients (**Figure 4K-L**). cRV patients had mostly increased expression of lysosome pathway transcripts, and many of these transcripts were reduced in dRV patients (Figure 4M). These data suggested our porcine data recapitulated many of the observed findings in humans, and that autophagy is impaired in multiple species of RVD.

### GP130 activation suppressed autophagy in iPSC-CMs

To define the direct effects of GP130 activation on cardiomyocyte autophagy, we treated human induced pluripotent stem cells differentiated into cardiomyocytes (iPSC-CMs) with the GP130 ligand, oncostatin M (OSM), or the autophagy inhibitor, bafilomycin (BAF), and quantified cellular area positive for light chain 3 (LC3) puncta. OSM significantly increased LC3 positive area (control: 8±2%, OSM: 17±9%, *p*-value: 0.0001), which was nearly identical to BAF (13±3%) (Figure 4N-O). In an independent experiment, we found SC144 prevented the accumulation of LC3 puncta in iPSC-CM treated with OSM but importantly not following BAF exposure (OSM: 3±1% BAF: 5±2% *p*-value: 0.0001) (Figure 4P-Q). Thus, these *in vitro* data supported our *in vivo* findings that GP130 signaling modulated RV cardiomyocyte autophagy, and these alterations were rescued by SC144 treatment only when GP130 signaling was engaged.

### GP130 inhibition normalized mitochondrial dimensions, cristae morphology, and attenuated metabolic pathway protein dysregulation

To probe how dysregulated autophagy impacted RV cardiomyocyte mitochondrial biology, we examined mitochondria morphology and global mitochondrial protein regulation. Transmission electron microscopy analysis demonstrated an accumulation of circular mitochondria with disrupted cristae morphology in PAB-Veh RV cardiomyocytes, which SC144 mitigated. (Figure 5 A-C). Quantitative proteomics of RV mitochondrial enrichments delineated how these morphological alterations associated with mitochondrial protein regulation. Hierarchical cluster analysis (HCA) and partial least squares discriminant analysis (PLS-DA) revealed GP130 antagonism modulated the RV proteome signature as PAB-SC144 animals clustered with control. (Figure 5 D-E). Kyoto Encyclopedia of Genes and Genomes (KEGG) pathway analysis of the top 250 proteins important for differentiating the three groups identified in the PLS-DA how SC144 impacted mitochondrial function. Six of the top 20 pathways were related to metabolism, and oxidative phosphorylation was one of the most enriched pathways (Figure 5F). When we specifically probed oxidative phosphorylation, we found nearly all protein subunits of complexes I-V were downregulated in PAB-Veh, which SC144 combatted (Figure 5 G-H). Because fatty acids are the primary carbon source used for ATP generation in the heart^53^ and fatty acid oxidation was enriched in our KEGG analysis, we probed proteins in the mitochondrial beta-oxidation pathway. In PAB-Veh RVs, most fatty acid oxidation proteins were reduced, but treatment with SC144 maintained FAO protein abundances (Figure 5 I-J). Taken together, these data revealed PAB-Veh RV cardiomyocytes had altered mitochondrial morphology and metabolic protein regulation, which SC144 treatment mitigated. Importantly, these changes were congruent with our snRNA-seq data.

**Figure 5:**
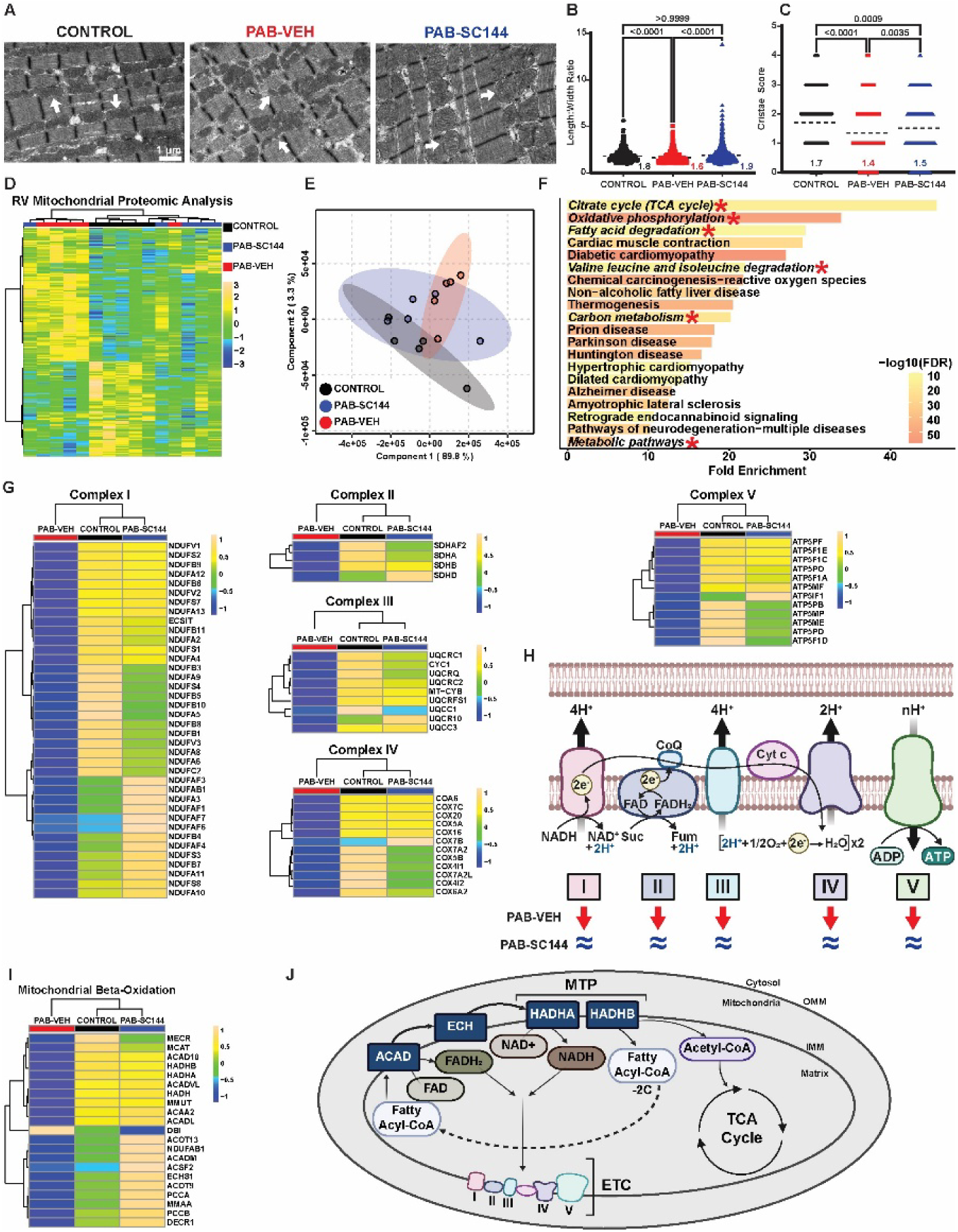
GP130 inhibition normalized mitochondrial dimensions, cristae morphology and mitigated dysregulation of oxidative phosphorylation and fatty acid metabolism proteins. (**A**) Representative electron micrograph demonstrating PAB-Veh mitochondria are smaller and more fragmented (white arrows). GP130 antagonism with SC144 normalized (**B**) mitochondrial shape assessed by length to width ratio (control: 1.8±0.7, PAB-Veh: 1.6±0.6, PAB-SC144: 1.9±0.9), and (**C**) mitigated changes in cristae abundance (control: 1.7±1, PAB-Veh: 1.4±1, PAB-SC144: 1.5±1). (**D**) Hierarchical cluster analysis and (**E**) partial least squares discriminant analysis of RV mitochondrial enrichment proteomics cluster PAB-SC144 clusters with control (**F**) Pathway analysis identified 6 of the top 20 pathways are related to metabolism (italicized, red star). (**G**) Proteins in complexes I-V of the electron transport chain were downregulated in PAB-Veh mitochondria, and normalized with SC144 treatment. (**H**) Schematic of complexes I-V of the electron transport chain. Red arrows (PAB-Veh) and blue equal signs (PAB-SC144) visualize changes compared to control abundances. (**I**) Proteins in the mitochondrial beta-oxidation pathway were reduced in PAB-Veh RVs, but not PAB-SC144 RVs. (**J**) Key proteins in the mitochondrial beta-oxidation pathway which produce cofactors for the electron transport chain and Acetyl-CoA for the TCA cycle. ACAD: acyl-coA dehydrogenase; ECH: enoyl-CoA hydratase; HADHA: hydroxyacyl-CoA dehydrogenase subunit A; HADHB: hydroxyacyl-CoA dehydrogenase subunit; MTP: mitochondrial trifunctional protein. Scale bars show relative changes between groups in **D**,**G**,**I**. Kruskal-Wallis ANOVA with Dunn post hoc test in **B**,**C**.

### PAB altered the RV lipidomic and metabolomic signature

To further evaluate derangements in RV metabolism, we performed a combined lipidomics and metabolomics analyses of RV free wall specimens. Lipidomics analysis of 937 lipids revealed changes in the overall composition of lipid species in the RV (Figure 6A). In particular, both PAB-Veh and PAB-SC144 RVs had a reduction in the total percentage of triacylglycerols (TAG) compared to control (control: 21%, PAB-Veh: 1%, PAB-SC144: 3%). This was accompanied by an increase in phosphatidylcholines (PC, control: 32%, PAB-Veh: 41%, PAB-SC144: 40%), and phosphatidylethanolamines (PE, control: 37%, PAB-Veh: 45%, PAB-SC144: 44%), the two most abundant phospholipids in cell membranes^54^. Hierarchical cluster analysis grouped PAB-SC144 and PAB-Veh together, suggesting GP130 antagonism minimally altered global lipid homeostasis (Figure 6B). Random forest classification identified triacylglycerol species as important lipids that differentiated the 3 groups (Figure 6C). We completed an enrichment analysis using Lipid Ontology (LION, Figure 6D)^44^ to further evaluate changes in RV lipid species in an unbiased manner. LION analysis revealed triacylglycerols, glycerolipids and fatty acids were the most enriched pathways. Similar to the lipidomics data, HCA of the top 200 metabolites with differential abundances clustered PAB-SC144 and PAB-Veh groups together (Figure 6E). Pathway analysis revealed amino acid metabolism pathways comprised many of the most enriched pathways when the three experimental groups were compared (Figure 6F).

**Figure 6:**
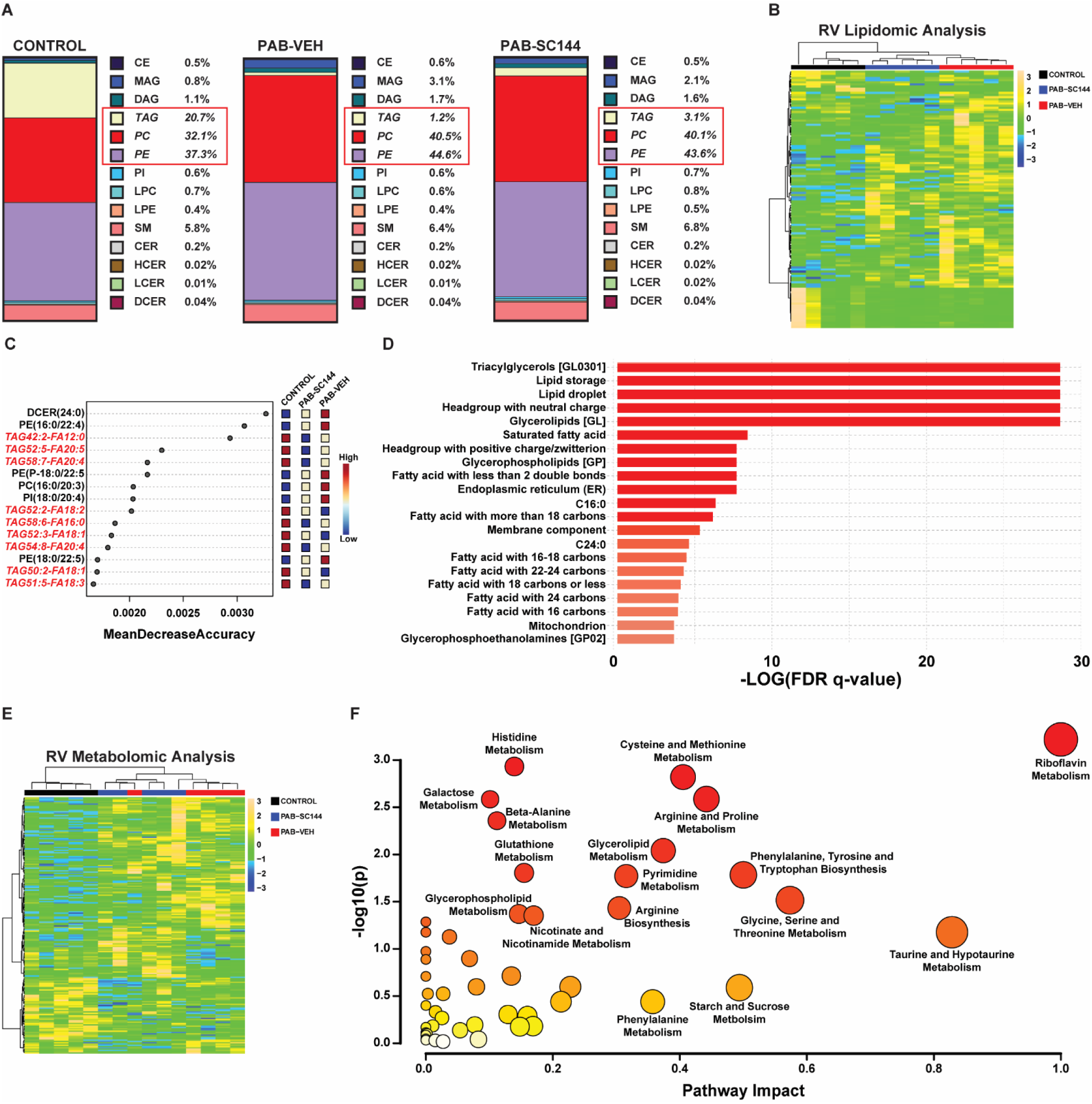
Combined lipidomics and metabolomics analyses identified altered RV metabolism. (**A**) PAB-Veh and PAB-SC144 had a reduction in the total percentage of triacylglycerols (TAG, control: 21%, PAB-Veh: 1%, PAB-SC144: 3%), and an increase in phosphatidylcholines (PC, control: 32%, PAB-Veh: 41%, PAB-SC144: 40%) and phosphatidylethanolamines (PE, control: 37%, PAB-Veh: 45%, PAB-SC144: 44%). (**B**) Hierarchical cluster analysis of lipidomics clustered PAB-Veh and PAB-SC144 groups together. (**C**) Random forest classification identified the top 15 lipids in distinguishing between experimental groups. 9 of 15 are TAG species. (**D**) Lipid Ontology enrichment analysis revealed triacylglycerols, glycerolipids, and fatty acids were the most enriched pathways. (**E**) Hierarchical cluster analysis of metabolomics data clustered PAB-Veh and PAB-SC144 groups together. (**F**) Pathway analysis of metabolomics data identified amino acid metabolism pathways account for many of the most enriched pathways. Scale bars show relative changes between samples in **B**,**E**

### SC144 counteracted changes in lipid/fatty acid metabolism

Next, we integrated our lipidomics and metabolomics data to assess how SC144 influenced RV fat metabolism. When we evaluated the abundances of individual TAG species, they were all reduced in both PAB-Veh and PAB-SC144 pigs (**Figure 7A-7B**) consistent with our relative abundance data. Similar numbers of diacylglycerols (DAG) were dysregulated between the three groups (Figure 7C). However, monoacylglycerols (MAG) were mostly increased in PAB-Veh pigs compared to control and PAB-SC144 pigs (Figure 7D), which suggested this step in fatty acid metabolism was uniquely impaired in PAB-Veh, but rescued by SC144. Then, we evaluated levels of free fatty acids and acylcarnitines. PAB-Veh RVs had reduced abundances of long chain fatty acids compared to controls, but these species were mostly increased in PAB-SC144 (Figure 7E). However, medium chain fatty acids and acylcarnitine were increased in PAB-Veh pigs, but normalized to control levels by SC144 (Figure 7F-H). Importantly, the concentration of carnitine, the cofactor required for the generation of acylcarnitines for mitochondrial import, was not significantly different between PAB-Veh and control pigs in both RV tissue and serum (**Supplemental Figure 5**). These data pointed to derangements in fatty acid metabolism in porcine RVD at three distinct steps: at MAG homeostasis, medium chain fatty acid regulation, and acylcarnitine degradation. Importantly, all three derangements were corrected with SC144.

**Figure 7:**
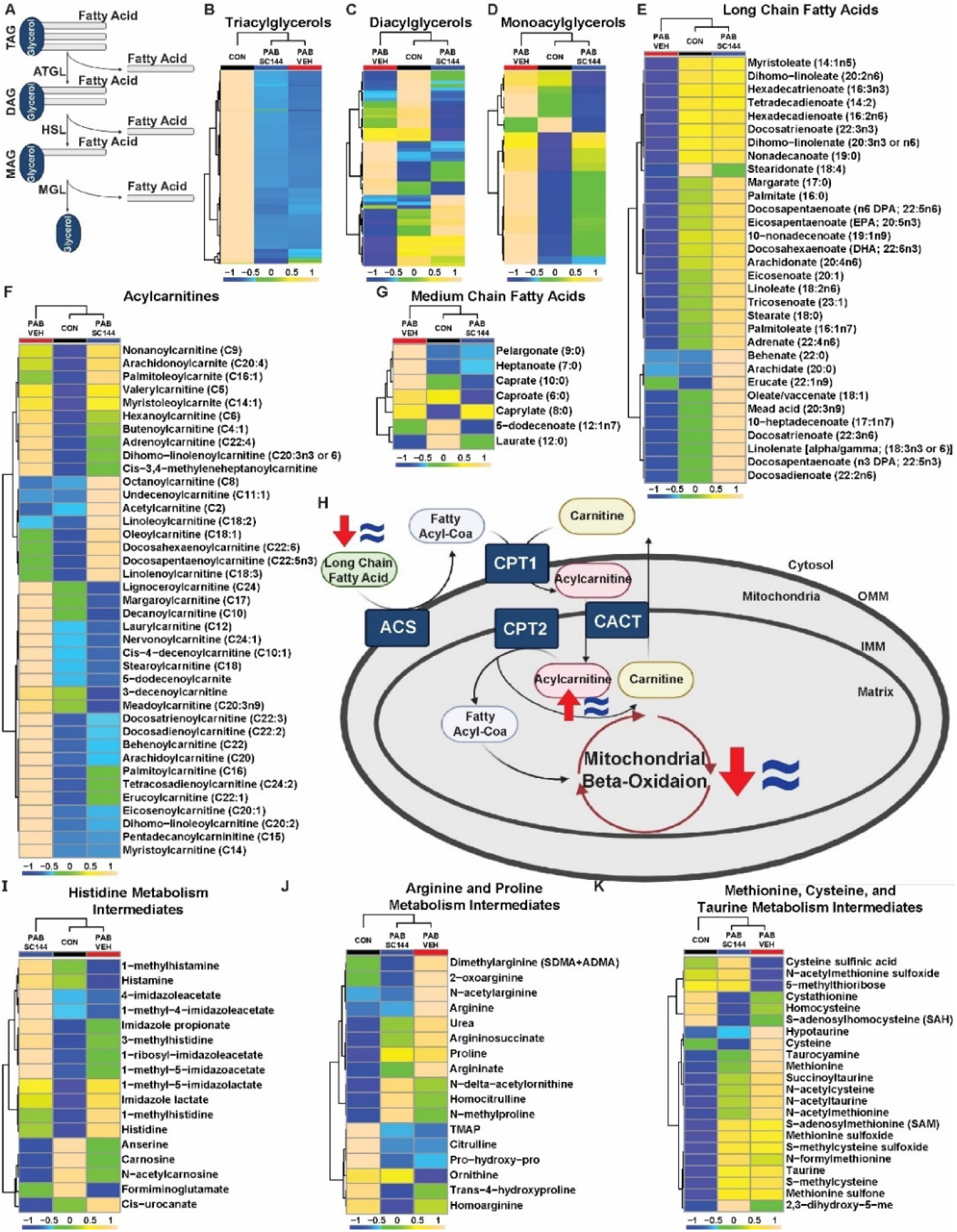
SC144 counteracted changes in lipid and fatty acid metabolism, but did not markedly alter amino acid metabolism. (**A**) Central steps in the breakdown of triacylglycerols into diacylglycerols and monoacylglycerols. ATGL: adipose triacylglycerol lipase; HSL: hormone sensitive lipase; MGL: monoacylglycerol lipase (**B**) Abundances of triacylglycerols are reduced in PAB-Veh and PAB-SC144. (**C**) Diacylglycerols were similar between all groups. (**D**) Abundances of monoacylglycerols are elevated in PAB-Veh RVs. (**E**) Long chain fatty acid species are decreased in PAB-Veh RVs compared to control and PAB-SC144. (**F**) Abundances of medium chain fatty acids are increased in PAB-Veh pigs. (**G**) Most acylcarnitines are increased in PAB-Veh RVs compared to control and PAB-SC144 RVs. (**H**) Key proteins and metabolites of the acylcarnitine shuttle in mitochondria. Red arrows (PAB-Veh) and blue equal signs (PAB-SC144) visualize changes in these metabolites compared to control. A carnitine molecule is exchanged for the CoA of a fatty acyl-CoA, and the subsequent acylcarnitine is transported into the mitochondrial matrix. The carnitine molecule is removed, and the fatty acyl-CoA undergoes beta-oxidation. ACS: acyl-CoA synthetase; CPT1: carnitine palmitoyltransferase I; CACT: carnitine acylcarnitine translocase; CPT2: carnitine palmitoyltransferase II. (**I**) PAB-Veh and PAB-SC144 RVs had similar abundances of metabolites in the histidine metabolism pathway. (**J**) Many metabolites in the arginine and proline, and (**K**) methionine, cysteine, and taurine metabolism pathways were increased in PAB-Veh and PAB-SC144 RVs. Scale bars show relative changes between groups in **B**-**G**, **I-K**

### SC144 did not markedly alter amino acid metabolism

In addition to fatty acid metabolism, our metabolomics pathway analysis revealed amino acid metabolism pathways were disrupted by PAB. Nearly all amino acid metabolites had similar abundances in PAB-SC144 and PAB-Veh RVs, and they frequently accumulated in RV tissue. The most divergent amino acid profiling was with histidine metabolism, as most intermediates were higher in PAB-Veh but reduced in PAB-SC144 (Figure 7I). However, metabolites in the arginine and proline, and the methionine, cysteine, and taurine metabolism pathways, were similarly dysregulated in PAB-Veh and PAB-SC144 RVs (Figure 7J-K). These data revealed GP130 antagonism primarily affected fat metabolism, as SC144 treatment minimally altered amino acid metabolites.

### PAB did not reduce lipid/fatty acid availability

Finally, we performed serum lipidomic and metabolomic analyses to determine if the RV changes we observed were potentially due to altered substrate delivery. TAG, DAG, and MAG species were mostly elevated in PAB-Veh serum compared to PAB-SC144 and control (**Supplemental Figure 6A-6C**). Both long and medium chain fatty acids were increased in the serum of PAB-Veh pigs (**Supplemental Figure 6D-E**). Serum acylcarnitines were similarly elevated in both PAB-Veh and PAB-SC144 pigs compared to control (**Supplemental Figure 6F**). Taken together, these data suggested that the derangements in RV fat metabolism were not likely a manifestation of reduced systemic lipid or fatty acid availability. Serum metabolomics identified many metabolites in the histidine, arginine and proline, and methionine, cysteine and taurine pathways were elevated in both PAB-Veh and PAB-SC144 (**Supplemental Figure 6G-6I)**, a pattern that mostly matched the RV changes. In conclusion, these data suggested the alterations in fatty acid metabolites were not due to systemic alterations and thus were RV-specific. However, the alterations in RV amino acids metabolites appeared to reflect systemic dysregulation.

## Discussion

In this study, we demonstrate GP130 antagonism significantly increases RVEF, mitigates RV cardiomyocyte hypertrophy, and rebalances the RV cellular landscape in PAB pigs (Figure 8). Data from snRNAseq, phosphoproteomics, electron microscopy, and mitochondrial proteomics reveals SC144 reduces cardiomyocyte mTORC1 signaling, which counteracts the downregulation of autophagy/lysosomal proteins, a finding observed in human RVD, restores mitochondrial morphology and cristae architecture, and normalizes the abundances of proteins responsible for fatty acid combustion and nearly all subunits of the electron transport chain. We corroborate these *in vivo* findings with *in vitro* experiments as GP130 stimulation impairs autophagy homeostasis in iPSC-CM, a phenotype that SC144 rescues. Furthermore, the combination of global metabolomics and lipidomics analyses reveals SC144 specifically enhances fatty acid metabolism. Moreover, SC144 reduces the abundance of macrophages and lymphocytes and blocks a pro-inflammatory genetic signature in both cell types. In summary, these data demonstrate GP130 antagonism improves RV function by restoring mitochondrial fatty acid metabolism through the rebalancing of mTORC1 associated autophagy and anti-inflammatory effects on lymphocytes and macrophages.

**Figure 8:**
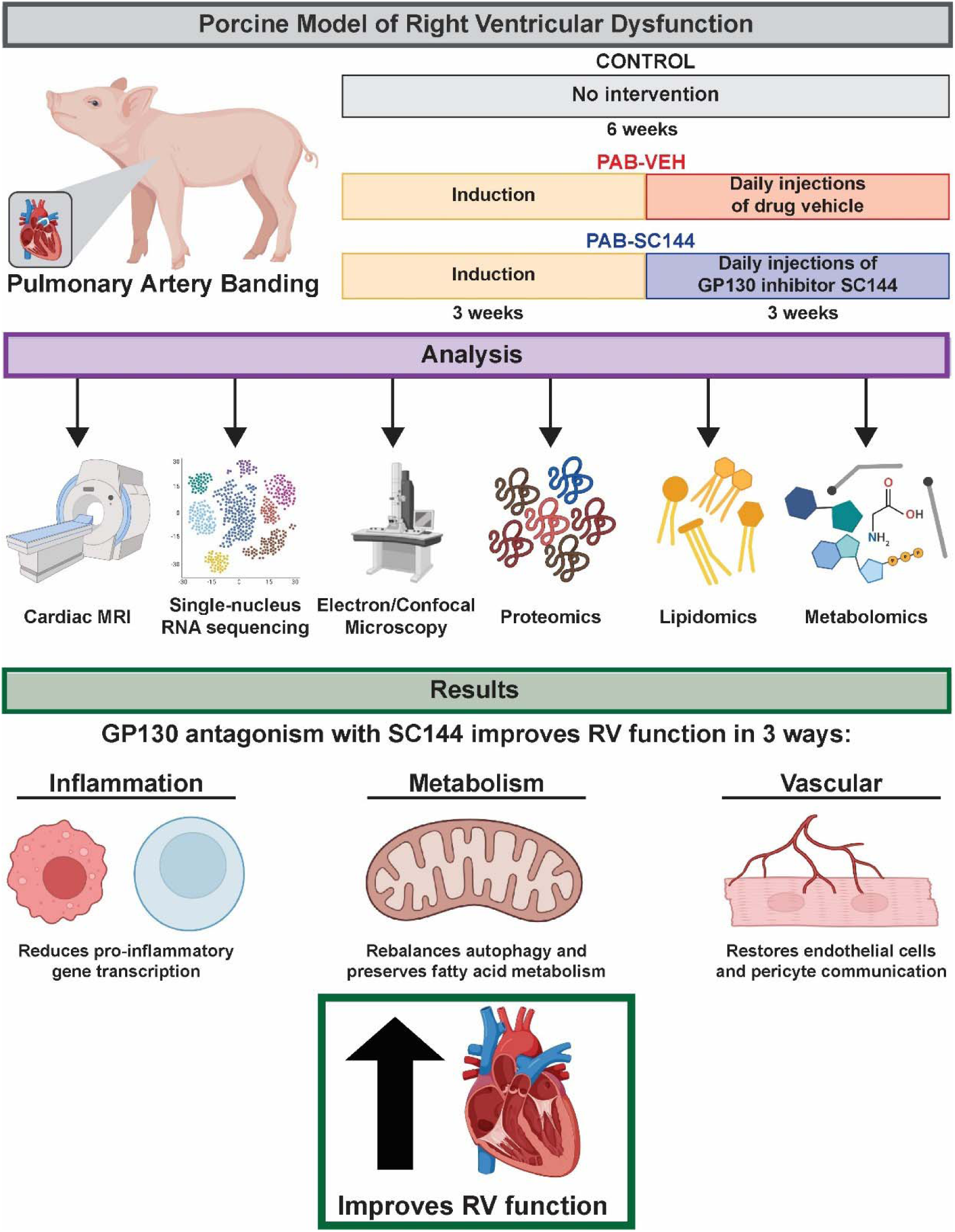
Proposed mechanism demonstrating GP130 antagonism with SC144 improves RV function. SC144 reduces the abundance of macrophages and lymphocytes and blocks pro-inflammatory gene transcription in both cell types, rebalances autophagy and preserves fatty acid metabolism, and restores endothelial cell and pericyte communication.

Our finding that GP130 antagonism enhances RV function via metabolic mechanisms is congruent with data from multiple independent laboratories showing GP130 adversely modulates metabolism and GP130 signaling exerts cardiotoxic effects. First, whole body knockout of the GP130 ligand IL11 or anti-IL11 therapy in aged mice improves metabolism in multiple tissues via a proposed mTORC1-associated mechanism^14^. In cardiac biology, expression of a constitutively active GP130 mutant heightens post myocardial infarction mortality due to LV rupture^55^. In addition, cardiac-specific knockout of the endogenous GP130 inhibitor, suppressor of cytokine signaling 3 (SOCS3), exacerbates cardiac dysfunction following pressure overload^56^. Moreover, SOCS3 knockout mice spontaneously develop dilated cardiomyopathy with high mortality^56, 57^. Furthermore, SOCS3 knockout animals have altered mitophagy and display downregulation of genes involved in mitochondrial metabolic pathways, including oxidative phosphorylation and fatty acid metabolism^58^. In conclusion, there are data from multiple species with different physiological insults demonstrating GP130 signaling has pathogenic effects via impaired metabolism.

We nominate derangements in the mTORC1-autophagy-fatty acid metabolism axis as the underlying cause of mitochondrial structural and functional deficits in RV cardiomyocytes. Although the role of autophagy in RVD is understudied, autophagy in human left heart failure and animal models of left ventricular dysfunction is the subject of many investigations. Notably, failing human hearts have reduced autophagic clearance^59^, which supports our findings that this pathway has pathogenic implications. We propose restoration of autophagy homeostasis normalizes mitochondrial fatty acid oxidation, which underlies some of the cardioprotective effects of SC144. There are data to support this hypothesis as augmentation of fatty acid metabolism imparts beneficial effects in RVD. Increasing fatty acid oxidation counteracts RV dysfunction in rodent models^38, 60, 61^ ^62^, and a phase II clinical trial shows metformin, an activator of fatty acid oxidation, improves RV fractional area change in patients with PAH^63^. On the contrary, there are also robust data from rodents and humans demonstrating suppressing fatty acid oxidation is beneficial in RVD. Inhibition of fatty acid oxidation inhibition with ranolazine and trimetazidine improves RV function in rodents^64^, and ranolazine increases RV ejection fraction in patients with pulmonary hypertension associated RVD^65^. Beyond altering bioenergetics, suppressing fatty acid oxidation promotes cardiac regeneration in adult mice^66^, which could underlie some of the benefits ranolazine and trimetazine impart in both rodents and humans.

Our lipidomics analysis is the first large-scale evaluation of lipid changes in a large animal model of RV dysfunction, and thus provides additional insights into mechanisms underlying RVD. Triacylglycerol (TAG), monoacylglycerol (MAG), phosphatidylethanolamine (PE), and phosphatidylcholine (PC) lipid species are the most robust changes when comparing control to diseased animals. TAG are predominately storage vessels for fatty acids for eventual metabolism^67^, and thus the reduction in TAG likely reflects an increase in energy demands following PAB. This agrees with our findings that TAG species are reduced in the RVs of monocrotaline rats^68^. The accumulation of MAG in RVD may have deleterious consequences as MAG can disrupt Golgi structure in intestinal cells, which could alter Golgi-based protein trafficking and regulation^69^. Alterations in PC and PE homeostasis may also have important implications. Disrupted PC/PE ratio reduces mitochondrial respiration, ATP synthesis, and increases mitochondrial fragmentation^54^, which could be another reason underlying the mitochondrial dysfunction observed in RVD. Furthermore, polyunsaturated PE are the lipid species that promote ferroptosis^70^, a lipid-dependent form of cell death, and suppressing ferroptosis improves RV function in PAB-pigs^20^. Because PCs and PEs are similarly increased in PAB-Veh and PAB-SC144 pigs, the above-mentioned mechanisms may explain why GP130 antagonism does not completely restore mitochondrial metabolism or RV function. Surprisingly, lipotoxic ceramides are not elevated in PAB-Veh RVs (**Supplemental Figure 7**). This finding is contrary to data from rodents^38, 61^ and humans^71^ demonstrating ceramide levels are elevated in RVD. Thus, our porcine model exhibits less lipotoxicity than rodents and humans with RVD. In summary, our lipidomics analysis provides new insight into RVD and identifies potential mechanisms underlying SC144’s inability to completely rescue the RV phenotype.

In our snRNAseq analysis, we define alterations in endothelial cells and pericytes in RVD, which likely contribute to RV vascular dysfunction. Porcine RVD results in greater abundances of endothelial cells and pericytes. While GP130 antagonism minimally alters pericyte abundance, it actually increases the abundance of endothelial cells (Figure 2D-F). Pathway analysis of differentially expressed genes in endothelial cells suggests their repair mechanisms, but not their replicative capacity, are impaired in RVD (**Supplemental Figure 8A**). Apelin signaling^72^, a reparative pathway in endothelial cells, is reduced in PAB endothelial cells while MAPK, a proliferative pathway, is enriched. These data point to a potential imbalance between repair and replication, which may help explain why the RV microvasculature morphology, but not density, is altered in RVD^73^. The RV vasculature is also dependent on pericytes, cells that support angiogenesis and ensure proper vascular function^74^. Interestingly, GP130 signaling is already implicated in pericyte biology as IL6 stimulates vessel sprouting, but inadequate pericyte coverage in *ex vivo* rodent aortas^73^, showing GP130 activation inhibits optimal pericyte function. In cardiac biology, a reduction in pericyte abundance reduces vascular integrity and increases capillary leakage following myocardial infarction^74^. Our data suggest a related phenomenon may be occurring in RVD because the Ras pathway, which promotes proper pericyte-endothelial cell interactions during angiogenesis^75^, is reduced in PAB-Veh pericytes (**Supplemental Figure 8B**). Furthermore, when comparing predicted communication between PAB-Veh and PAB-SC144 endothelial cells and pericytes, CellChat analysis identifies 3 ligand-receptor pairs unique to PAB-SC144 (**Supplemental Figure 8C**). The platelet-derived growth factor D to platelet-derived growth factor receptor beta (PDGFD-PDGFRB) is only predicted from PAB-SC144 endothelial cells to pericytes. PDGFD, when released by endothelial cells, increases pericyte recruitment^76^ and therefore PAB-Veh endothelial cells likely have reduced pericyte coverage. In addition, the laminin-α2 to integrin subunit α6/integrin β1 (LAMA2-ITGA6+ITGB1) and LAMA2-ITGA1+ITGB1 are only identified from PAB-SC144 pericytes. LAMA2 signaling is necessary for proper vessel formation and is more highly expressed in mature pericytes^77^. In summary, suppressing GP130 activation may restore defects in the RV vasculature via modulation of endothelial cell and pericyte biology, which may be another mechanism by which SC144 augments RV function.

Our animal data supports the translational potential of GP130 antagonism in RVD and potentially provides insights into why modulating this pathway failed in the past. Certainly, GP130 antagonism needs to be considered carefully as it may have immunosuppressive effects, but Olamkicept, a soluble GP130 receptor ligand trap^78^, shows a promising safety profile and clinical benefit in inflammatory bowel disease patients^78, 79^. We hypothesize that GP130 antagonism enhances RV function beyond simply inhibition of IL6 as our snRNA-sequencing data show the leukemia inhibitory factor receptor (LIFR), IL11 receptor alpha (IL11Ra), and oncostatin M receptors (OSMR) have the highest expression in the RV. Notably, the IL6 receptor (IL6R) is minimally expressed in the porcine RV (**Supplemental Figure 9**). This observation may explain why IL6 antagonism with Tocilzumab does not reduce NT pro-BNP levels in PAH patients^80^. Importantly, we demonstrate GP130 antagonism exerts pleiotropic effects via inflammatory, metabolic, and vascular changes, which may improve its translatability as data suggests all these pathogenic mechanisms contribute to human RVD^11, 12, 73^.

## Limitations

Our study has important limitations that we acknowledge. First, we used young, castrated male piglets and thus our findings may not be translatable to adults, but examination of human data from other laboratories suggests GP130 signaling is activated in RVD^11, 12^. We chose to first evaluate the effects of GP130 antagonism in a porcine model using male pigs because in rodent models of RVD males have a more severe phenotype^81^, and thus females may not have the same response as males. However, human data probed for this study (Figure 4E-J) included male and female patients^11^. Due to housing constraints, pigs needed to be castrated which could alter GP130 signaling. However, intact and castrated male pigs exhibit similar increases in plasma IL6 levels after lipopolysaccharide challenge^82^, which suggests this pathway is still intact in castrated animals.

Our snRNA-sequencing data show a higher proportion of cardiomyocytes than analyses of adult hearts^28^. However pediatric hearts have a greater relative percentage of cardiomyocytes^83^, and perhaps our model may better recapitulate pediatric RVD than adult disease. GP130 signaling is necessary for cardiomyocyte development and proliferation postnatally as GP130 inhibition reduces cardiomyocyte size and function in murine LVs. Surprisingly, RV cardiomyocytes are unaffected by GP130 antagonism and there is no RV dysfunction with GP130 inhibition^83^, which suggests the RV may be a better target for GP130 signaling than the LV. Our cMRI analysis does not identify any LV defects with SC144 (**Supplemental Table 1**), and in fact the augmentation of RV function mitigates minor LV dysfunction (**Supplemental Table 1**). We identify changes in the electron transport chain and mitochondrial beta-oxidation pathways that are divergent from one of our previous porcine publications^18^, but consistent with another^20^. This finding may be due to the difference in degree of RV dysfunction between the two studies. In a previous cohort of banded pigs that exhibit less RV dysfunction (RVEF of 38%^18^), there are increases in ETC and FAO proteins, but in pigs with more severe RVD (RVEF of 30%^20^) we see reductions in ETC and FAO proteins^20^. We did not observe as robust a fibrotic response in our porcine model as observed in monocrotaline rats. Castration may be a contributor as mouse studies show castration can mitigate RV fibrosis^84^. Finally, our previous work suggests GP130 signaling stimulates pathogenic microtubule stabilization, which contributes to RV pathology^9, 85^. There is a reduction in RV tubulin subunit protein abundances with SC144, but the upregulation of tubulin subunits in PAB-Veh (**Supplemental Figure 10**) is not as marked as we see in rodents^9, 60^.

---

**Supplemental Table 1:** Complete cMRI data. Data are presented as mean ± standard deviation or median and [interquartile range]. Subscript “i” indicates value indexed to final pig weight.

|  | **Control (n=5)** | **PAB-VEH (n=5)** | **PAB-SC144 (n=5)** |
| --- | --- | --- | --- |
| **RA EDV<sub>i</sub> (mL/kg)** | 1.1 [0.9-1.1] | 1.5 [1.4-1.9] | 1.4 [0.9-2.3] |
| **RA ESV<sub>i</sub> (mL/kg)** | 0.4 [0.4-0.5] | 0.9 [0.8-1.1] | 0.7 [0.5-1.6] |
| **RA SV<sub>i</sub> (mL/kg)** | 0.6 ± 0.1 | 0.6 ± 0.2 | 0.6 ± 0.2 |
| **RA EF (%)** | 60 [54-63] | 35 [34-46] | 46 [33-53] |
| **RA SV/ESV** | 1.5 [1.2-1.7] | 0.5 [0.5-0.9] | 0.9 [0.5-1.1] |
| **RV EDV<sub>i</sub> (mL/kg)** | 2.4 ± 0.5 | 4.9 ± 1.8 | 3.6 ± 1.3 |
| **RV ESV<sub>i</sub> (mL/kg)** | 1.0 ± .2 | 3.9 ± 1.9 | 2.3 ± 1.0 |
| **RV SV<sub>i</sub> (mL/kg)** | 1.4 ± 0.3 | 1.0 ± 0.2 | 1.3 ± 0.5 |
| **RVEF (%)** | 59 ± 4 | 22 ± 10 | 37 ± 6 |
| **RV Mass<sub>i</sub> (g/kg)** | 0.8 ± 0.2 | 1.8 ± 0.4 | 1.5 ± 0.4 |
| **RV SV/ESV** | 1.5 [1.3-1.7] | 0.3 [0.1-0.5] | 0.5 [0.5-0.8] |
| **LV EDV<sub>i</sub> (mL/kg)** | 3.3 ± 0.5 | 3.0 ± 0.2 | 2.4 ± 0.3 |
| **LV ESV<sub>i</sub> (mL/kg)** | 1.7 ± 0.4 | 1.8 ± 0.4 | 1.3 ± 0.3 |
| **LV SV<sub>i</sub> (mL/kg)** | 1.6 ± 0.2 | 1.1 ± 0.3 | 1.1 ± 0.1 |
| **LV EF** | 48.2 ± 5 | 39 ± 11 | 47 ± 6 |
| **RVEDV/LVEDV** | 0.7 [0.7-0.8] | 1.4 [1.2-2.4] | 1.3 [1.1-1.9] |
| **TR Severity** | NA | 1.8 ± 0.4 | 1.5 ± 0.4 |
| **Proximal PA CSA** | NA | 2.8 ± 0.6 | 2.6 ± 0.4 |
| **PA-Band CSA** | NA | 0.3 ± 0.1 | 0.4 ± 0.1 |
| **% Stenosis** | NA | 89 ± 2 | 86 ± 5 |

**Supplemental Figure 1:**
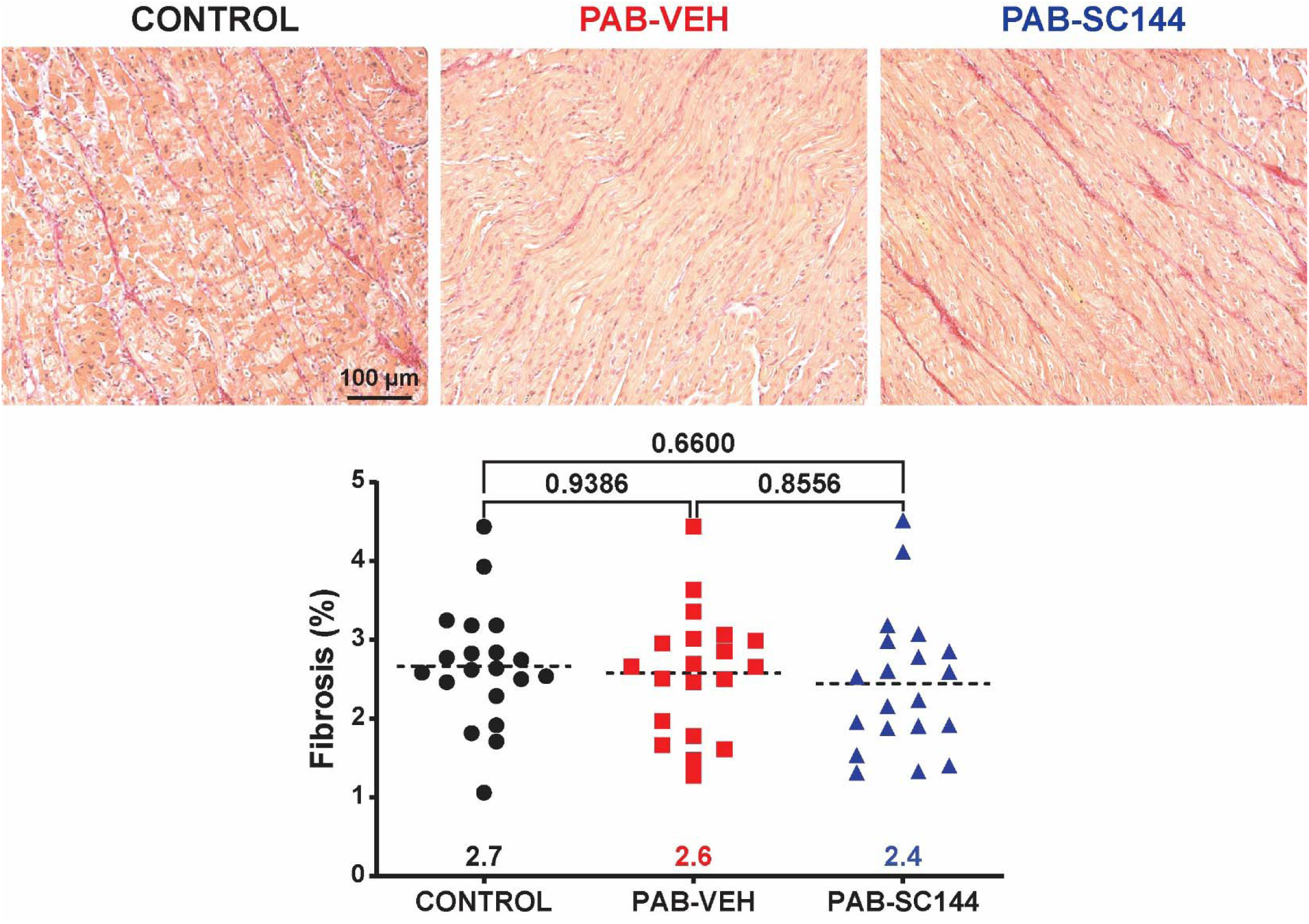
Neither PAB nor SC144significantly affected fibrosis. Representative microscopy images of RV sections stained with picrosirius red to evaluate fibrosis. Fibrosis was similar between all three groups (control: 2.7±0.7%, PAB-Veh 2.6±0.8%, PAB-SC144: 2.4±0.9%)

**Supplemental Figure 2:**
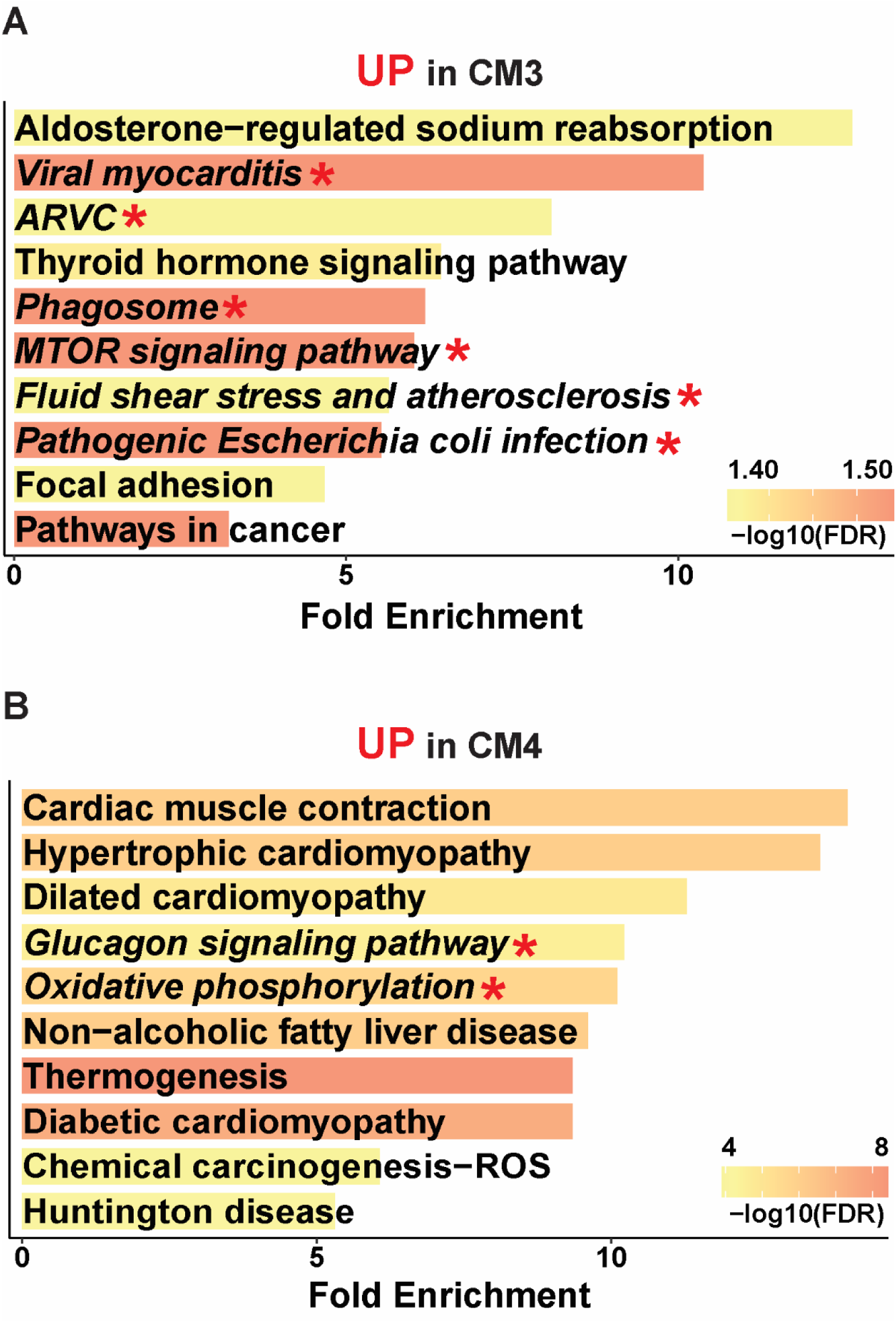
SC144 counteracted metabolic gene expression dysregulation in cardiomyocytes. (**A**) Pathways enriched in cardiomyocyte subgroup 3 (CM3), found mostly in PAB-Veh RVs, were related to inflammation and mTOR signaling (italics, red star). **(B)** Metabolic pathways, including oxidative phosphorylation (italics, red star), were elevated in CM4, which were identified at similar proportions in control and PAB-SC144.

**Supplemental Figure 3:**
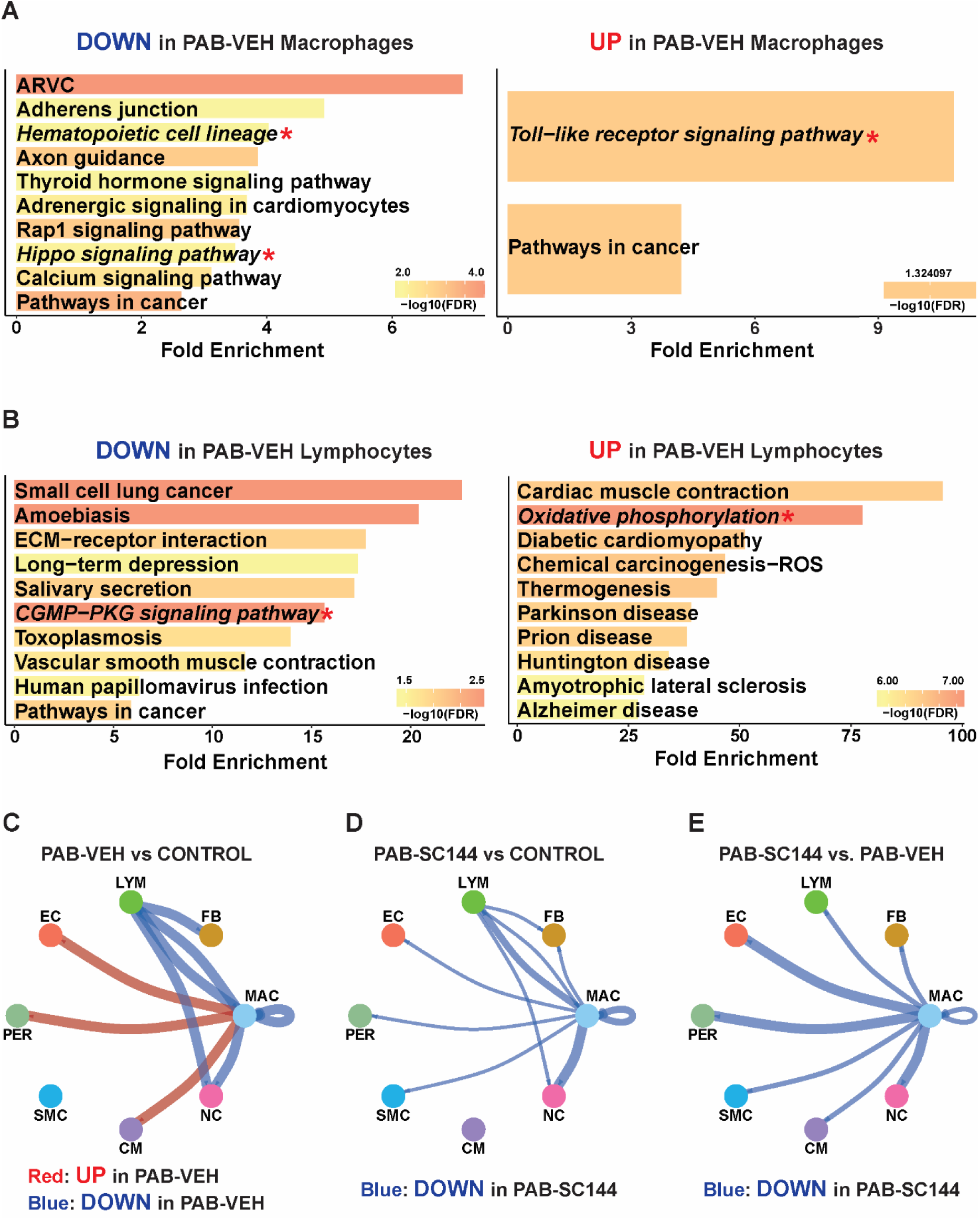
Pro-inflammatory pathways and predicted communication from immune cells are increased in PAB-Veh immune cells. Compared to control, (**A**) pathways that regulate macrophage differentiation, survival and proliferation (italics, red star) were suppressed in PAB-Veh, and the pro-inflammatory pathway toll-like receptor signaling was enriched in PAB-Veh macrophages. (**B**) Anti-inflammatory pathways were reduced in PAB-Veh lymphocytes relative to control, and oxidative phosphorylation was increased. CellChat communication analysis showing (**C**) predicted communication from macrophages to endothelial cells, pericytes, and cardiomyocytes was higher (red) in PAB-Veh macrophages compared to control, but (**D**) reduced (blue) in PAB-SC144 immune cells compared to control. (**E**) Compared to PAB-Veh, PAB-SC144 immune cells had reduced predicted signaling.

**Supplemental Table 2:**
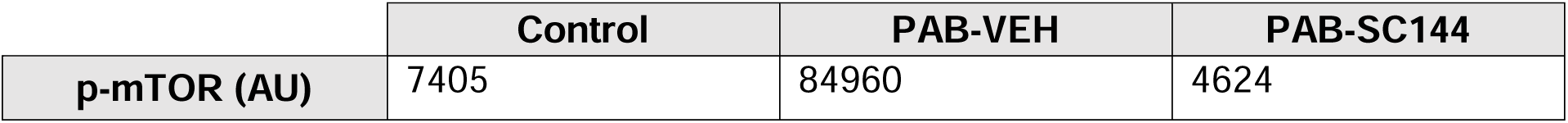
Grouped abundances of phosphorylated mTOR (p-mTOR). p-mTOR was elevated in PAB-Veh, but normalized by SC144.

**Supplemental Figure 4:**
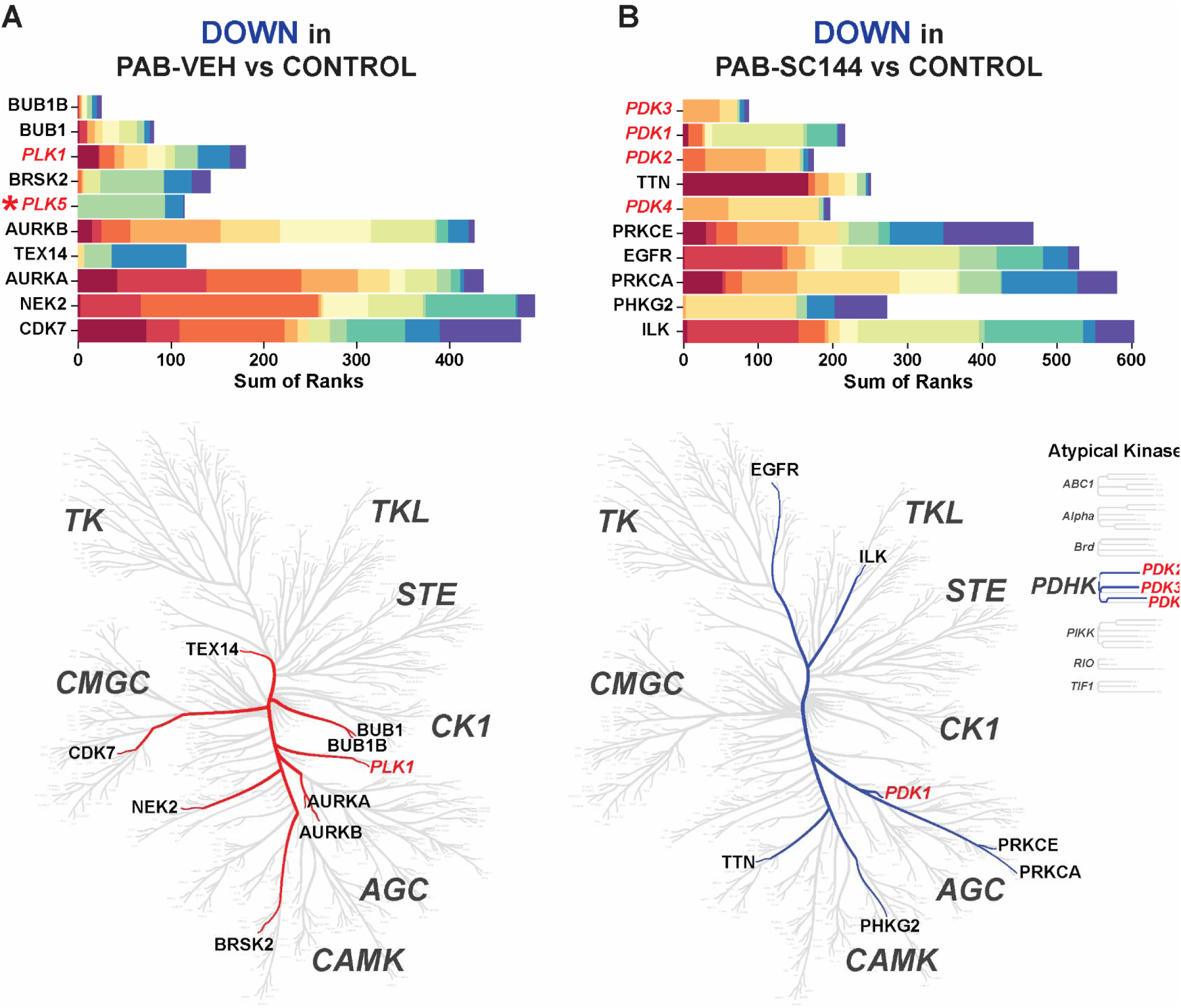
GP130 antagonism with SC144 is predicted to alter metabolic kinases. (**A**) Several polo like kinases (PLK, red italics) which antagonize mTORC1 were estimated to be diminished in PAB-Veh compared to control. PLK5 was not in the Coral Database and is therefore not visualized on the kinome map (red star). (**B**) Multiple pyruvate dehydrogenase kinases (PDK, red italics) which support glucose utilization over fatty acid oxidation were predicted to be reduced in PAB-SC144 compared to control.

**Supplemental Figure 5:**
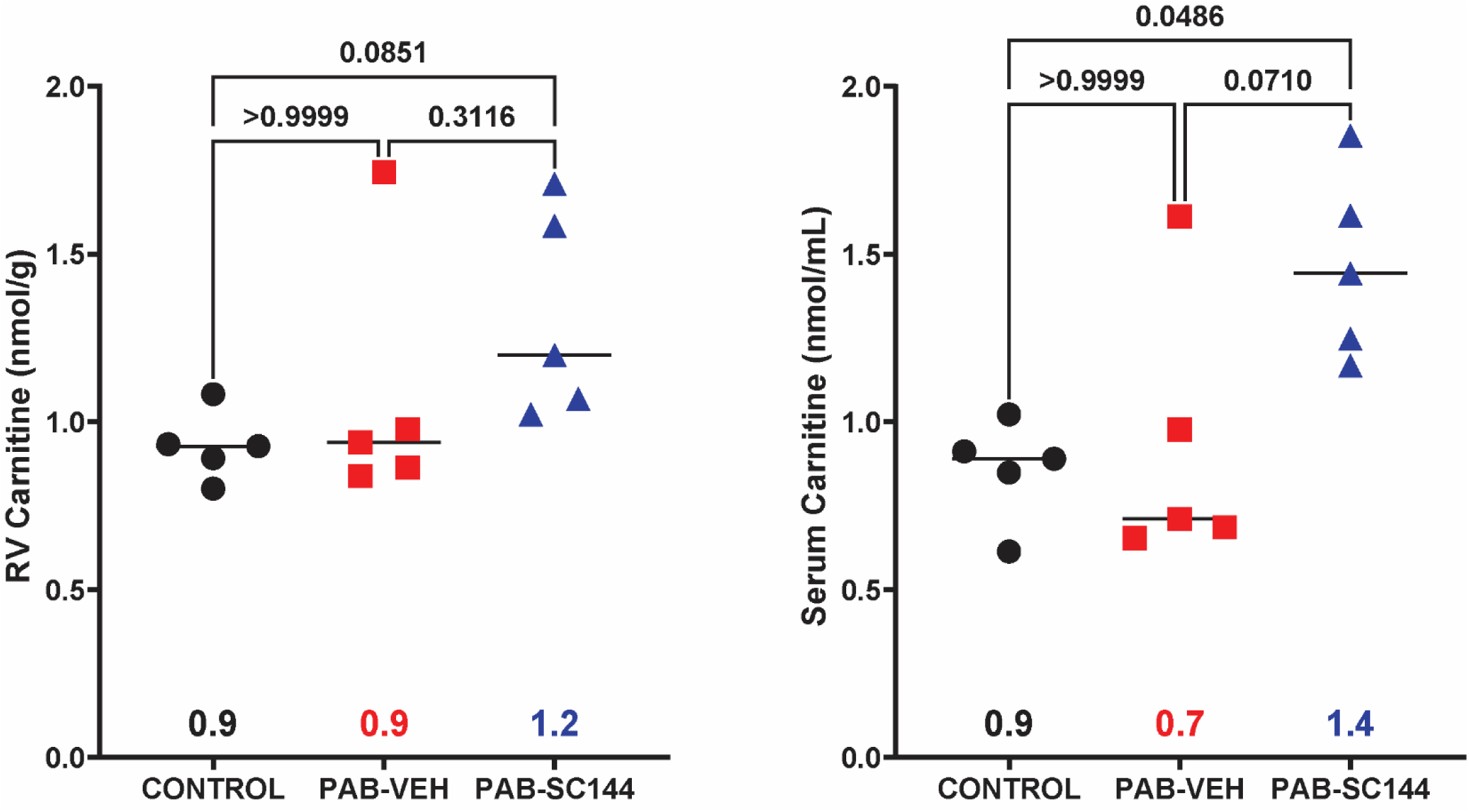
PAB did not affect RV or serum carnitine levels. RV (0.9, [0.8-1.0]) and serum (0.9, [0.7-1.0]) carnitine levels are similar between all 3 groups, but serum carnitine was slightly elevated in PAB-SC144 pigs. Kruskal-Wallis ANOVA with Dunn post hoc test used.

**Supplemental Figure 6:**
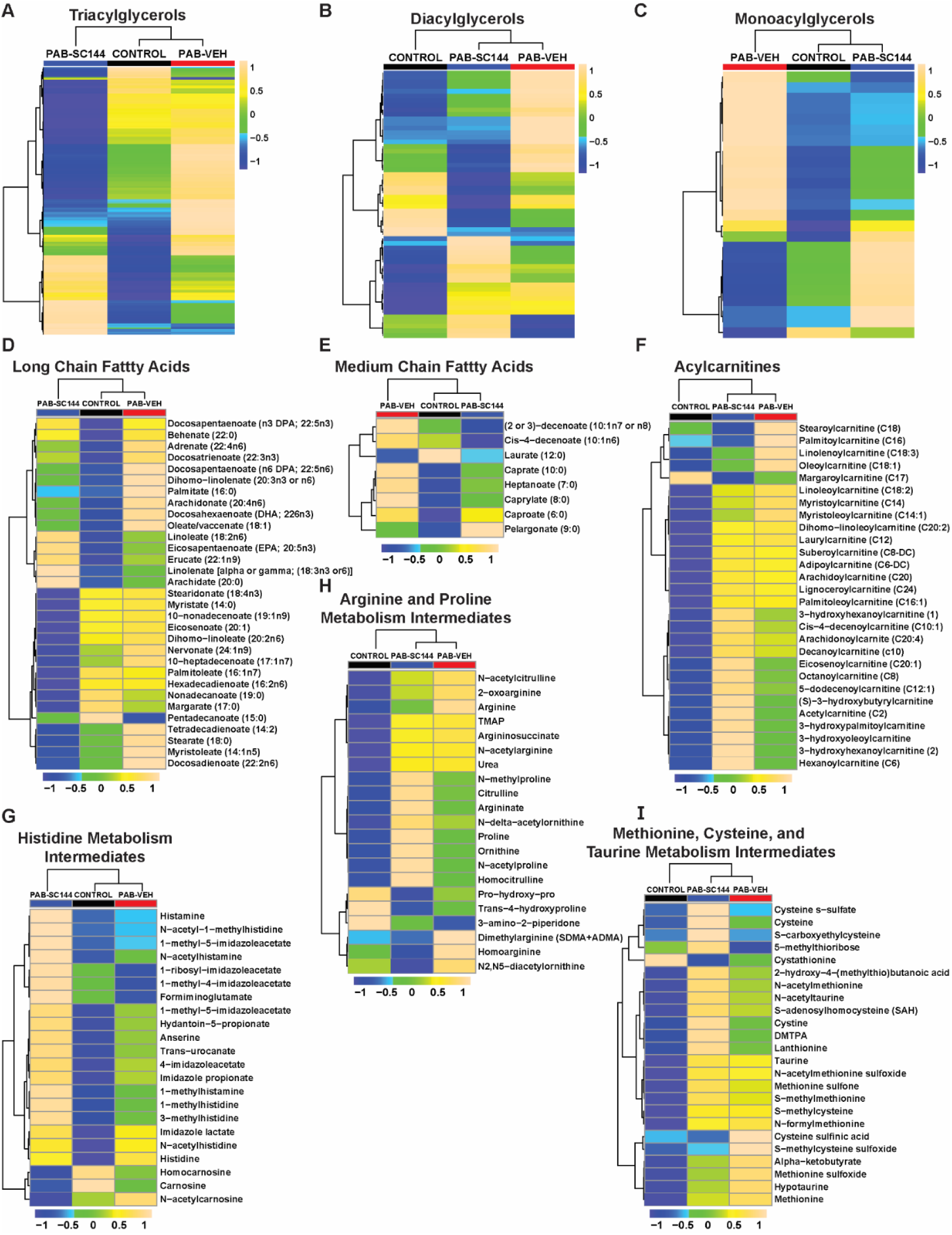
Serum lipidomics and metabolomics reveal changes in RV fat metabolism were not likely due to reduced systemic lipid or fatty acid availability. (**A**) TAG, (**B**) DAG, and (**C**) MAG species were mostly elevated in PAB-Veh serum compared to control and PAB-SC144. Serum (**D**) long and (**E**) medium chain fatty acids were increased in PAB-Veh, and acylcarnitines were similarly elevated in PAB-SC144 and control serum. Many amino acid metabolism intermediates in the (**G**) histidine, (**H**) arginine and proline, and (**I**) methionine, cysteine and taurine pathways were increased in both PAB-Veh and PAB-SC144. Scale bars show relative changes between groups.

**Supplemental Figure 7:**
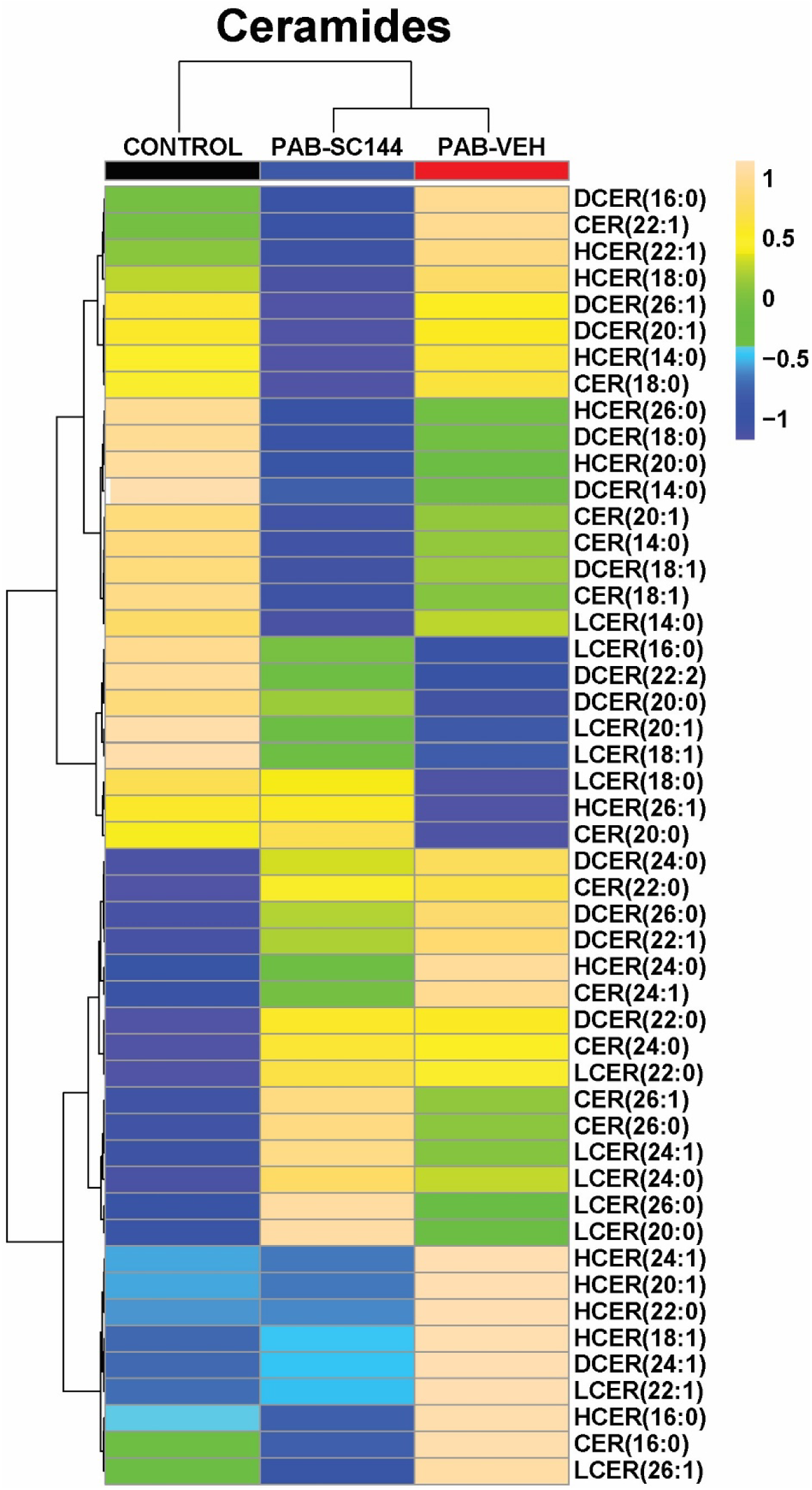
Ceramide species are not elevated in PAB-Veh RVs compared to control or PAB-SC144. Scale bar shows relative changes between groups

**Supplemental Figure 8:**
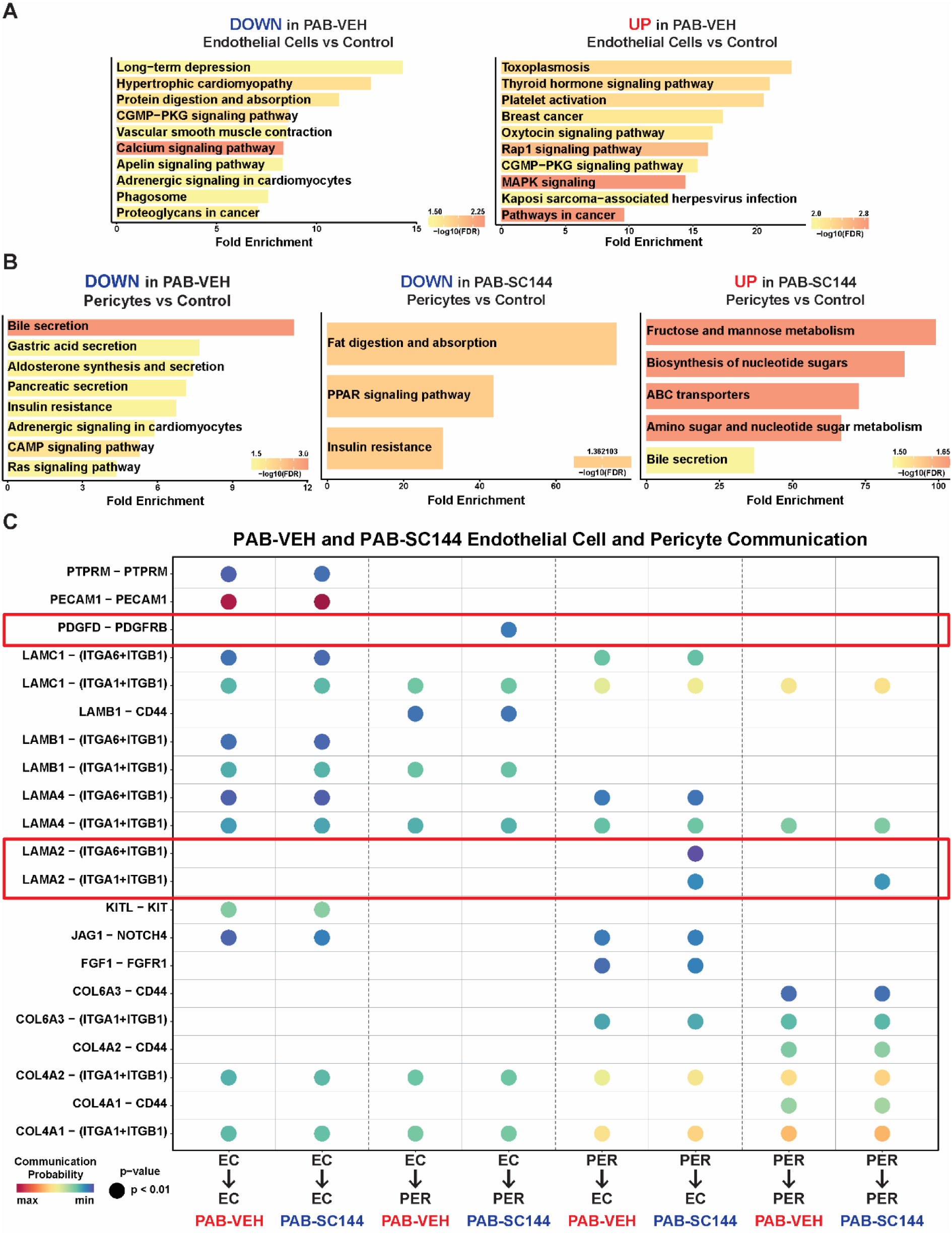
GP130 inhibition modulates endothelial cell and pericyte biology. (**A**) Pathway analysis of PAB-Veh DEGs compared to control identify endothelial cell repair mechanisms may be reduced, but replication is not altered. (**B**) Pathways that promote proper pericyte-endothelial cell interactions during angiogenesis are reduced in PAB-Veh pericytes. (**C**) CellChat analysis identified 3 ligand receptor pairs (red boxes) only enriched in PAB-SC144 vascular cells, not PAB-Veh that contribute to endothelial cell-pericyte stability.

**Supplemental Figure 9:**
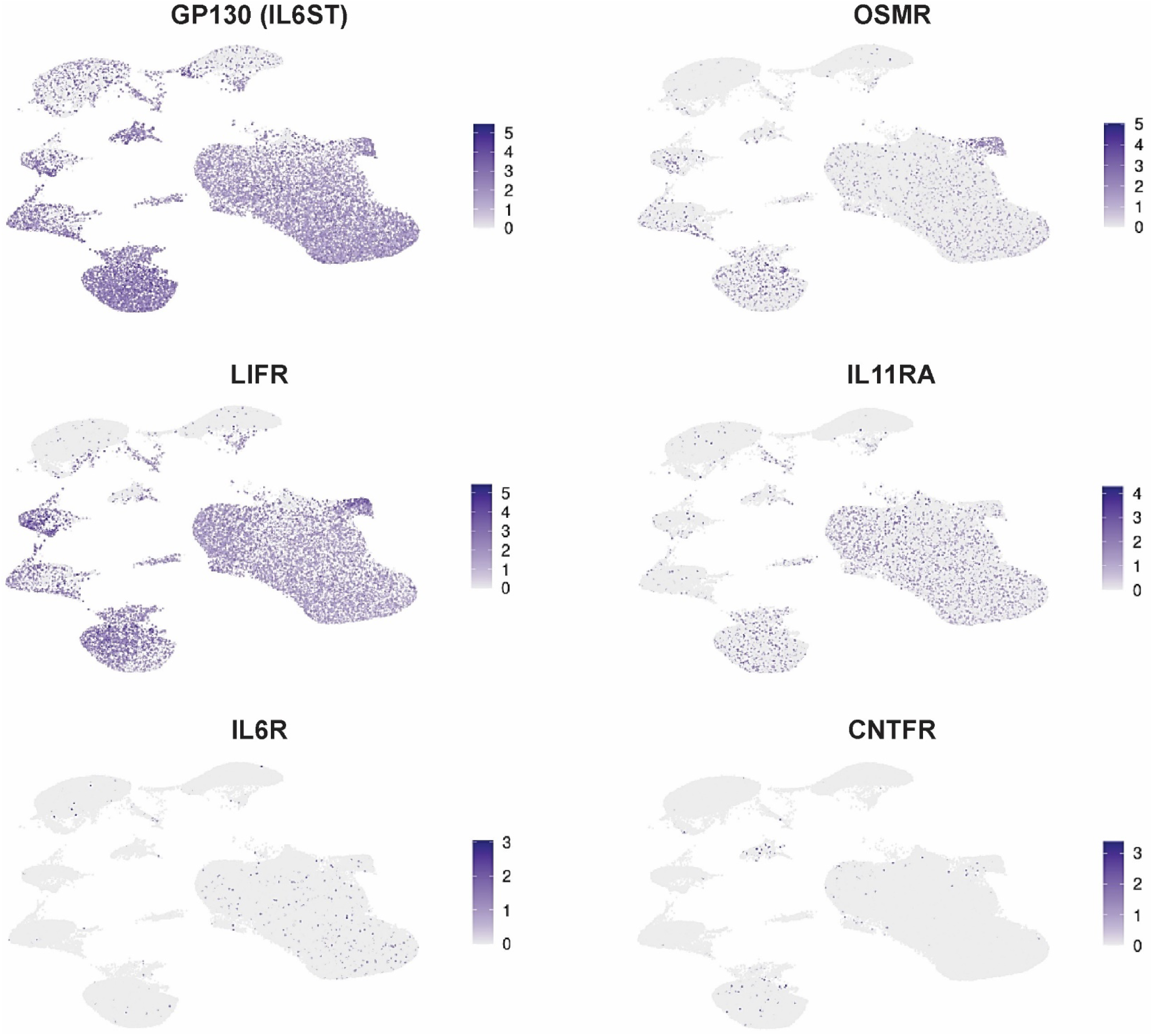
GP130 receptor expression across all cell types. Besides GP130, the leukemia inhibitory factor receptor (LIFR), IL11 receptor (IL11RA), and oncostatin M receptor (OSMR) had the highest expression. The IL6 receptor (IL6R) and ciliary neurotrophic factor receptor (CNTFR) had lower expression in the RV.

**Supplemental Figure 10:**
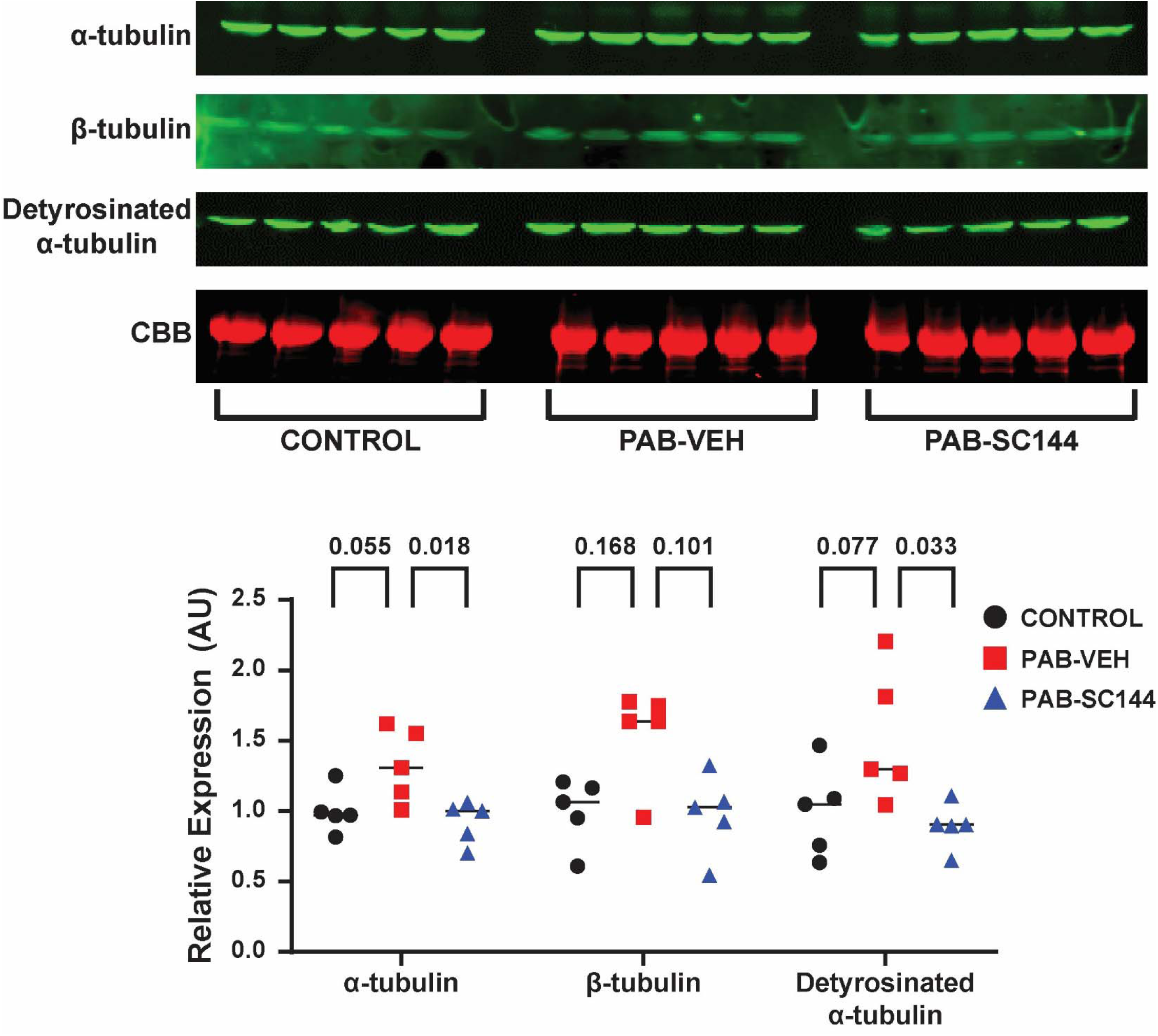
SC144 normalized RV tubulin abundances, but PAB did not cause a significant increase in these proteins. Representative western blots and quantification of alpha(α)-, beta(β)-, and detyrosinated α-tubulin.

